# Evolution of *Mycobacterium tuberculosis* transcription regulation is associated with increased transmission and drug resistance

**DOI:** 10.1101/2025.05.01.651750

**Authors:** Peter H Culviner, Abigail M Frey, Qingyun Liu, Dang Thi Minh Ha, Phan Vuong Khac Thai, Do Dang Anh Thu, Nguyen Le Quang, Roger Calderon, Leonid Lecca, Maxine Caws, Sarah J Dunstan, Megan B Murray, Nguyen Thuy Thuong Thuong, Sarah M Fortune

## Abstract

*Mycobacterium tuberculosis* (Mtb) has co-evolved with humans for thousands of years causing variation in virulence, transmissibility, and disease phenotypes. To identify bacterial contributors to phenotypic diversity, we developed new RNA-seq and phylogenomic tools to capture hundreds of Mtb isolate transcriptomes, link transcriptional and genetic variation, and find associations between variants and epidemiologic traits. Across 274 Mtb clinical isolates, we uncovered unexpected diversity in expression of virulence genes which we linked to known and previously unrecognized regulators. Surprisingly, we found that many isolates harbor variants associated with decreased expression of EsxA (Esat6) and EsxB (Cfp10), which are virulence effectors, dominant T cell antigens, and immunodiagnostic targets. Across >55,000 isolates, these variants associate with increased transmissibility, especially in drug resistant Mtb strains. Our data suggest expression of key Mtb virulence genes is evolving across isolates in part to optimize fitness under drug pressure, with sobering implications for immunodiagnostics and next-generation vaccines.

## Introduction

All pathogens face the continuous challenge to survive and transmit while avoiding clearance by host immunity or drug treatment. While many pathogens stay ahead in this arms race through rapid genetic diversification, *Mycobacterium tuberculosis* (Mtb) is highly genomically conserved. Mtb does not engage in horizontal gene exchange and has a lower mutation rate than many other pathogens^1,2^. Even strains from different lineages that diverged thousands of years ago only differ by a few thousand bases. Despite the organism’s relative genetic monomorphism, experimental and epidemiologic studies suggest that Mtb strains differ in disease manifestations, rates of relapse, and transmissibility^3,4^.

We recently analyzed clinical Mtb strain genomes to identify sites of diversifying selection occurring within the course of single human infections, which we postulated would include functionally significant variants driving drug or host immune escape^5^. In addition to canonical drug resistance genes, we found that many targets of within host selection are transcription factors, some of which have known roles in virulence including *phoPR*, *devR*, and *whiB6*. Interestingly, genes from three two-component signal transduction systems (*phoR, kdpE* and *narS*) were recently also identified as sites of temporally episodic diversifying selection, suggesting regulatory selection may aid in adaptation to changing host dynamics^6^. Across life there is evidence regulators play a role in evolution: prokaryotic regulators are less conserved than their regulons and in eukaryotes regulatory variation is already a well appreciated mechanism of diversification^7,8^. We hypothesized that regulatory variants alter the expression of key bacterial pathways and contribute to Mtb adaptation and phenotypic variation.

Though there are tens of thousands publicly available of Mtb isolate genomes, the largest transcriptomic studies only include tens of strains^9,10^. We reasoned that broader analysis of Mtb transcriptomes might identify adaptive evolution of gene expression and allow us to predict its causal variants. To do this, we developed new methods to capture and analyze the transcriptomes from hundreds of Mtb strains^9,10^. We built new tools for high throughput microbial RNA-Seq including innovations for standardized growth and lysis, and a library workflow for non-polyadenylated RNA, which we term Multiplex Taq-depleted Bacterial (MTB) RNA-Seq that reduces the cost of library preparation to ∼$5 per sample. With this platform, we conducted RNA-Seq on 274 Mtb clinical isolates from two clinical cohorts spanning phenotypically drug sensitive strains from the 3 most prevalent Mtb lineages (Figure 1A)^11,12^.

**Figure 1.**
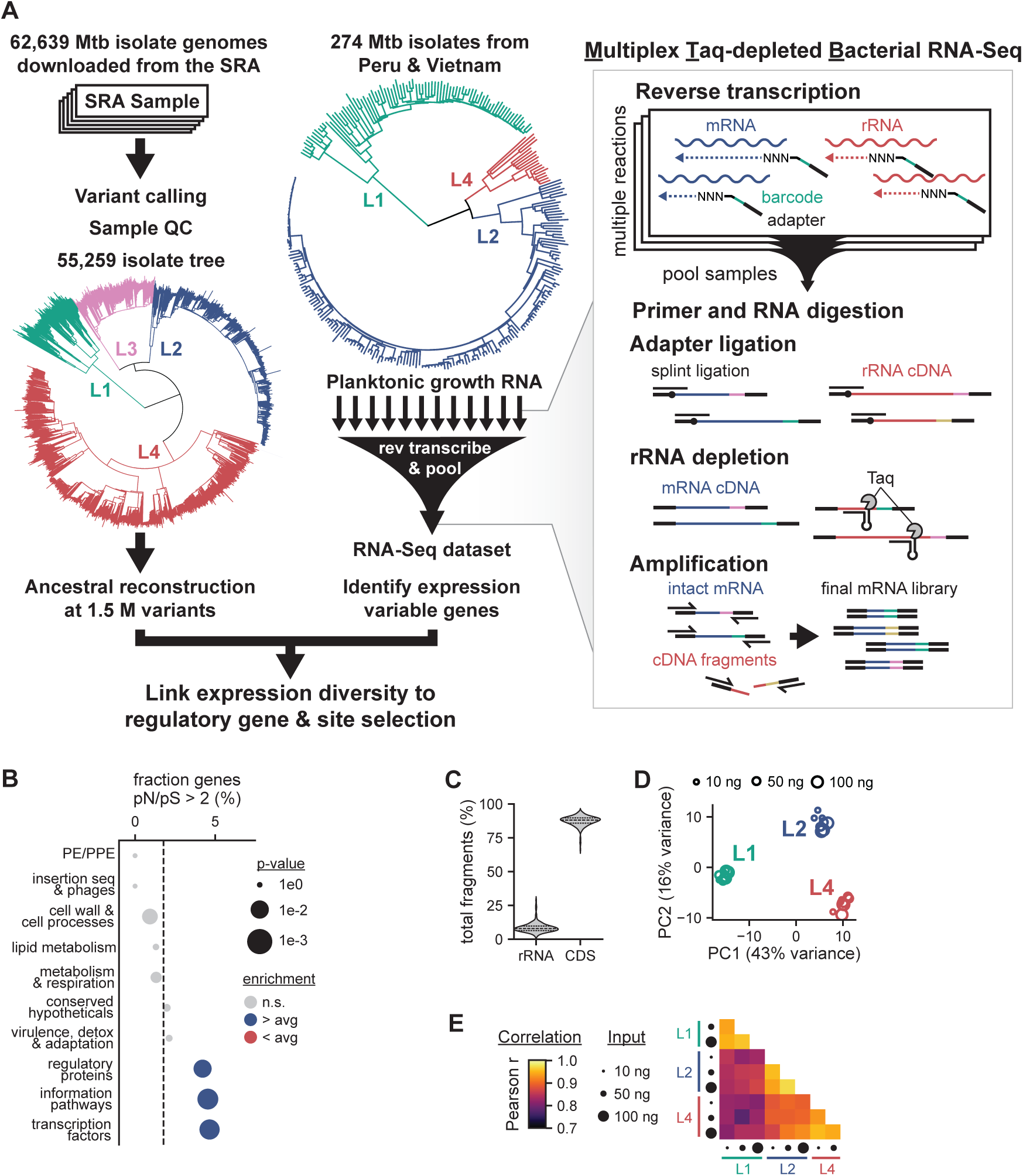
Multiplex Transcriptomics of Mtb Clinical Isolates. **(A)** Schematic workflow of (left) variant calling and ancestral reconstruction pipeline and (right) high throughput transcriptomic pipeline. Phylogenetic trees are shown with 4 major Mtb lineages colored. **(B)** Enrichment analysis of genes under diversifying selection and Mtb gene categories. Dots sizes reflect the −log_10_ p-value of 2-sided Fisher exact test with statistical significance shown by color. **(C)** Violin plots of percentage of total fragments falling in rRNA and coding regions from the libraries presented in this study. Mean and quartiles shown as dotted lines. **(D)** Principal component analysis of 27 validation libraries using different barcodes and 3 different concentrations of RNA from lineage 1, 2, and 4 samples. The log_2_ expression of the top 25% of well-expressed genes sorted by standard deviation were used. (**E)** Heatmap of Pearson correlations between mean log_2_ expression of all well-expressed genes for all validation libraries. See also Figure S1.

Across these isolates, we found high expression diversity in genes essential for survival in host and known antigens, in contrast to the high rate of sequence conservation across Mtb T cell antigens themselves^13^. Most surprisingly, we identified recurrent evolution of decreased expression in the *PE35-PPE68-esxBA* operon, which encodes critical virulence effectors secreted by the ESX-1 secretion system. *EsxBA* and ESX-1 more generally are essential for Mtb virulence and loss of ESX-1 is the primary attenuating deletion in the vaccine strain BCG^14^. Using genome wide association approaches, we find decreases in *PE35-PPE68-esxBA* expression are linked to prevalent genetic variants in the *whiB6* transcriptional regulator. Leveraging global genome sequencing data, we then show that *whiB6* variants are themselves associated with transmission success, especially in drug resistant strains. These results demonstrate that genetically conserved virulence pathways can be affected by adaptive regulatory selection and suggest a possible virulence–transmission tradeoff with implications for pathogen evolution and vaccine design^15^.

## Results

### Transcription factors are targets of diversifying selection

We began by defining genes under selective pressure across global Mtb isolates, leveraging publicly available whole genome sequencing data from 55,259 Mtb clinical strains^5^. In our previous analyses, we analyzed genes evolving within individual patients, but here we assessed fixed variants and therefore captured both ancient and modern evolution. Variant calling, quality control, merger, and analysis is computationally impractical with this number of genomes. Thus, we built a library of computational pipelines parallelizing download, calling, storage, and comparison of isolate genomes using existing tools (e.g. *tbprofiler*, *mutect2*, *breseq*, *pastml*), enabling rapid analysis and quality control of millions of variant calls across tens of thousands of isolates from the short read archive^16–21^. We used our pipeline and *fasttree2* to build 14 lineage-specific subtrees and combined them into a “super tree” to conduct parsimony ancestral reconstruction on all 55,259 variants^22,23^. Because calling variants on this number of strains requires a high-performance computing cluster, the trees, variant calls, and ancestral reconstructions are provided as community resources.

We used the variant calls and ancestral reconstruction to quantify selective pressure across the genome by calculating the ratio of non-synonymous to synonymous population-wide polymorphisms (pN/pS ratio) for every gene (Figure S1A, Table S1)^24^. In agreement with prior studies, the distribution of pN/pS ratios was biased towards purifying selection (mean=0.69)^6^. We next searched for enrichment of genes under diversifying selection (pN/pS ratio greater than 2) in at least one major lineage across broad functional categories as defined by *mycobrowser* (Figure 1B)^25,26^. Information pathways, regulatory proteins, and transcription factors were identified as targets of diversifying selection, suggesting a possible evolutionary pressure to modify gene expression.

### Scaling Mtb transcriptomics to hundreds of phenotypically diverse clinical isolates

We then sought to directly measure the extent of transcriptional variation between Mtb isolates. There are few large-scale analyses of transcriptional diversity across bacterial strains due to the technical complexity and expense of capturing many strains at a uniform growth state, extracting RNA and performing rRNA depletion in the absence of poly-adenylation^27^. Mtb adds further challenges because of its slow growth, growth rate variation between clinical strains, and the difficulty of cell lysis.

To address these challenges, we developed new tools for inexpensive, parallelized transcriptional profiling of hundreds of Mtb strains (Figure 1A, right). We first developed a Mtb culturing and RNA processing pipeline that enabled growth at the scale of hundreds of strains. We sought to measure gene expression in standard culture medium at mid-exponential phase of growth (OD_600_ ∼0.4, Figure S1B). This is technically challenging (especially at a ∼1 ml volume) because clinical Mtb strains have highly variable lag phases and doubling times. We used a digital camera to image culture turbidity and estimate OD_600_ from a validated standard curve. At the desired OD_600_, we added paraformaldehyde directly to the media to rapidly halt transcriptome dynamics and sterilize cultures. Crosslinks were reversed before RNA extraction.

We also developed a new RNA library pipeline, MTB RNA-Seq, to facilitate scaled analysis. Sample-specific inline barcodes were integrated during reverse transcription, the first step of library generation, to enable pooling of 48 samples after cDNA synthesis (Figure 1A, right). Then we used a new rRNA depletion approach that operates on pooled, barcoded cDNA instead of RNA. Our method uses Taq polymerase’s endonuclease activity to cleave rRNA reads using a DNA guide complementary to the rRNA combined with a loop that mimics the endonuclease’s natural DNA junction substrate (Figure S1C)^28^. Like CRISPR-based depletion, cleavage of these substrates prevents their amplification in the final library, but without the requiring PAM sites or RNA guide synthesis^29^. We demonstrated that cleavage of a target oligo was >99% efficient for matching guides if the target was single-stranded (Figure S1D). A library of guides targeting abundant non-coding RNAs enriched coding reads in our libraries (Figure 1C). The complete library generation protocol generated replicable clustering of the same RNA across a range of inputs (Figure 1D) and Pearson correlations of gene expression between replicate samples were high (Figure 1E).

### Mtb’s transcriptome reflects both ancient divergence and modern variation

We used our transcriptional profiling platform to assess gene expression of 274 Mtb strains in biologic duplicate during *in vitro* growth in 7H9 medium. These strains were phenotypically antibiotic-susceptible isolates from patients in Vietnam and Peru^11,12^. To identify genes with a high biologic not technical variation, we measured the ratio of the biologic variation between isolates to the technical variation between replicates (Figure 2A, Table S2). Most genes were distributed around 1, suggesting strain-to-strain and technical variation were of a similar magnitude, but there was a long tail of genes with greater biologic than technical variation. Here we focused on genes at the top 5% of biologic to technical variation; also, we excluded genes that did not have a high magnitude of biologic variation (bottom 50%). Applying these stringent thresholds, we identified 151 genes with high between strain expression diversity.

**Figure 2.**
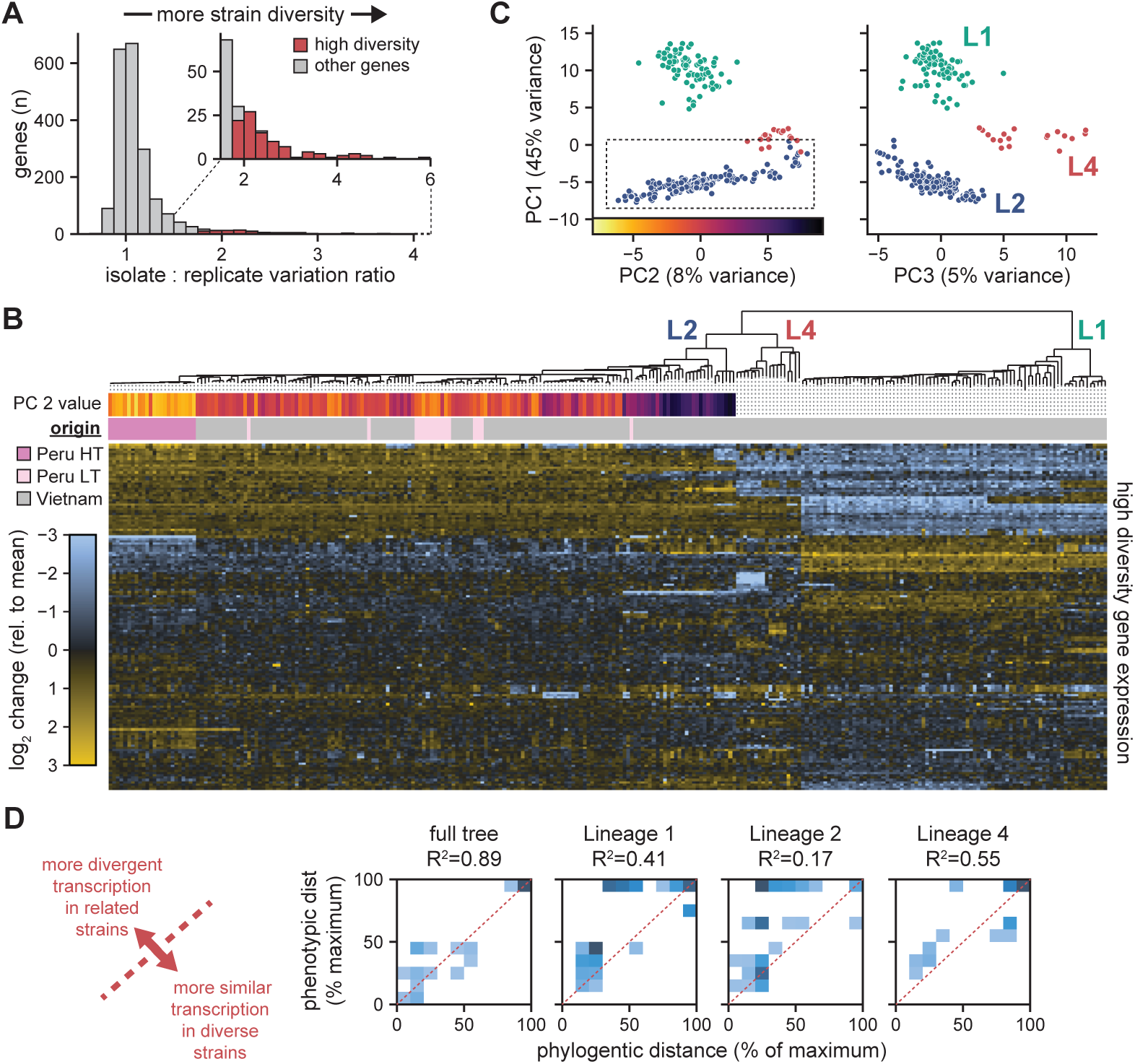
The Mtb transcriptome varies at ancient and modern timescales. **(A)** Histogram of the ratio of the mean between isolate standard deviation to the mean between strain standard deviation across all strains. We define high diversity genes as those that meet our expression threshold, are in the top 50% in mean between isolate standard deviation, and top 5% in isolate to replicate standard deviation. **(B)** Phylogenetic tree of Mtb isolates with transcriptional data with a heatmap of strain PC 2 value from 2C plotted against lineage 2 strains, source cohort, and a heatmap of mean strain gene expression for high diversity strains relative to the gene’s mean across all strains. **(C)** Scatterplots of principal component analysis separating strain (colored by lineage) transcriptional data using 151 high diversity genes as input. Component 1 is plotted against components 2 and 3 to highlight separation of lineage 2 (PC2) and separation of the lineages (PC3 x PC1). **(D)** Histograms of all strain pair phylogenetic patristic distances and phenotypic patristic distances calculated from Ward clustering high diversity gene expression. Clustering was conducted on the full tree as well as separately on each of the three major lineages. The R^2^ of the underlying scatters are shown. See also Figure S2.

To understand the genetic basis of this transcriptional diversity, we plotted the expression of the 151 high diversity genes against the strains’ phylogenetic tree (Figure 2B). Qualitatively, we observed breaks in gene expression patterns at every phylogenetic scale including lineage boundaries, smaller clades of closely related strains, and individual strains. High diversity gene expression robustly separated the three major Mtb lineages by principal component analysis (Figure 2C, right). PC1 separated the most phylogenetically distant strains (lineage 1 vs lineages 2 and 4) and accounted for a large fraction of the variance (45%). Lineage 2 strains had a large range of values in PC2 (Figure 2C, left) differentiating a recently expanded, highly transmissible Peruvian clade (Peru HT) from the more ancestral lineage 2 strains (Figure 2B, top)^12^. Deeper components provide additional discriminative power for clades diverging within the major lineages (Figures S2A, S2B). These results reflect transcriptional variation at ancient lineage boundaries but also indicate diversification of gene expression in more recently diverged clades.

To address this quantitatively, we compared the phylogenetic and phenotypic trees for all isolates together and separately for each lineage and plotted 2D histograms of the distances of every pair of strains. Across all strains, the phylogenetic and phenotypic distances closely matched (R^2^ = 0.89, Mantel test p-value < 1E-6; Figure 2D). For individual lineages, the correlations remained statistically significant (Mantel test p-value < 1E-6), but with lower R^2^ values of 0.41, 0.17, and 0.55 for lineages 1, 2, and 4, respectively. There were many pairwise comparisons with a much greater phenotypic distance than phylogenetic distance, many in lineage 2. These data suggest that transcriptome evolution does not occur at the same rate across the entire phylogenetic tree.

### Gene expression diversity is enriched in host-interactive genes and antigens

Genes with diverse expression belong to all the broad *mycobrowser* functional categories (Figure 3A, top). These include known virulence genes including *devRS*, involved in hypoxia regulation and persistence and known to vary transcriptionally^30^; *icl1*, involved in carbon metabolism and persistence^31^; and the *pks3-pks4-papA3-mmpL10* operon, involved in biosynthesis of immunomodulatory (poly-/di-) acyl trehalose (PAT/DAT; Figure S3A)^32^.

**Figure 3.**
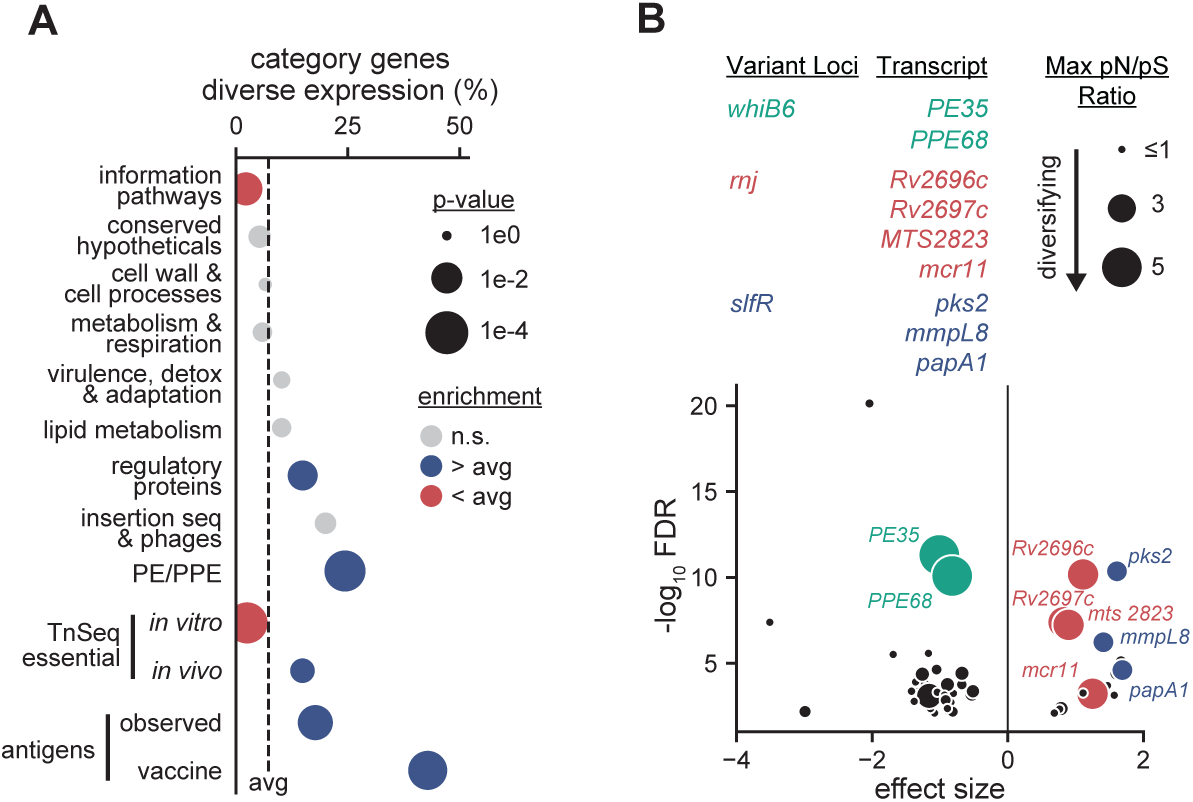
Mtb-host interactive processes have diverse expression. **(A)** Enrichment analysis of genes in the high expression diversity set vs functionally and experimentally defined gene categories from prior literature. Plotted as in 1B. **(B)** Volcano plot showing significant associations between transcripts enriched for expression outliers and the variable loci predictive of their expression. Four variant loci are color-coded, and their associated transcripts are labelled. Datapoints were sized according to the maximum pN/pS of the variant gene across L1-4 in Table S1. See also Figure S3.

Genes in information pathways—including core replication, transcription, and translation genes— were under-represented in the high variation gene set. Conversely, high variation genes were enriched for regulators and PE/PPE genes. PE/PPE genes are a group of proteins that saw a rapid expansion in Mtb^33^; though many of these proteins are uncharacterized, some are involved in nutrient uptake, and many are secreted and immunogenic. We used published TnSeq data to determine whether genes involved in homeostatic versus Mtb–host interaction vary in expression diversity. Expression diverse genes were under-represented in genes required for growth *in vitro*, while they were over-represented in genes required for survival and growth in a mouse model (Figure 3A, bottom)^34,35^.

We also considered the expression variation of Mtb antigens. Antigens from the Immune Epitope Database and vaccine-candidate antigens showed even stronger enrichment for expression diverse genes than *in vivo* essential genes^36–43^. In Mtb, T cell epitopes generally are conserved at a sequence level, perhaps because many play key roles during infection^6^, suggesting that transcriptional variation may provide a mechanism to tune immunogenicity or virulence without disrupting protein function.

### Linking expression diversity to genetic loci with genome wide association

We next linked high diversity gene expression to genetic variation using a genome wide association (GWAS) approach. As we have done previously, we assessed associations at a whole gene level, classifying strains as having a predicted functional variant in a gene or the reference state^44^. In Mtb, GWAS is complicated by complete genetic linkage. To focus on modern selection rather than associations driven by lineage-defining variants, we considered only variants that occur in less than 25% of strains and required gene loci to have at least 5 independent mutational events. We then used *pyseer* to associate expression diverse genes with genetic variants^45^.

We found 53 genomic associations between gene expression and genetic variants (Benjamini-Hochberg FDR α=0.01; Figure 3B, Table S3). These associations include potential *cis* regulatory relationships between adjacent genes that could be driven either by polar effects or autoregulatory loops, relationships between distant genes without known roles in gene regulation that could be driven by multi-site regulatory integration of changes in cellular state, and relationships between genes and known or predicted regulators. Three of these loci also had strong diversifying selection in one or more lineages: *rnj*, *Rv0042c* (*slfR*), and *whiB6*.

The Mtb ortholog of RNase J, *rnj*, was associated with increased abundance of the sRNAs *mcr11* and *mts2823,* and the operonic proteins *Rv2697c* and *Rv2696c* (Figure S3B). We previously identified RNase J as a target of recurrent loss of function mutations leading to multi-drug tolerance^46^. This study also identified *mcr11*, *mts2833* and *Rv2697c-Rv2696c* as differentially abundant in *rnj* mutants, building confidence in the results of our GWAS analysis.

### *Rv0042c* encodes *slfR*, a site of selection, and a novel regulator of sulfolipid-1 synthesis

Our GWAS also identified novel regulatory relationships which we next sought to validate. Specifically, variants in *Rv0042c* were associated differential expression of *pks2-papA1-mmpL8* (Figure 4A), which encode proteins involved in the synthesis and transport of sulfolipid-1 (SL-1) (Figure 4B)^47,48^. SL-1 is a virulence associated cell wall lipid recently implicated in cough induction through activation of nociceptive neurons^49,50^. Because 3 of 6 of the variants in *Rv0042c* encode pre-stops or frameshifts which were associated with increased *pks2-papA1-mmpL8* expression, we hypothesized that *Rv0042c*, hereafter referred to as *slfR*, encodes a repressor of SL-1 biosynthesis genes.

**Figure 4.**
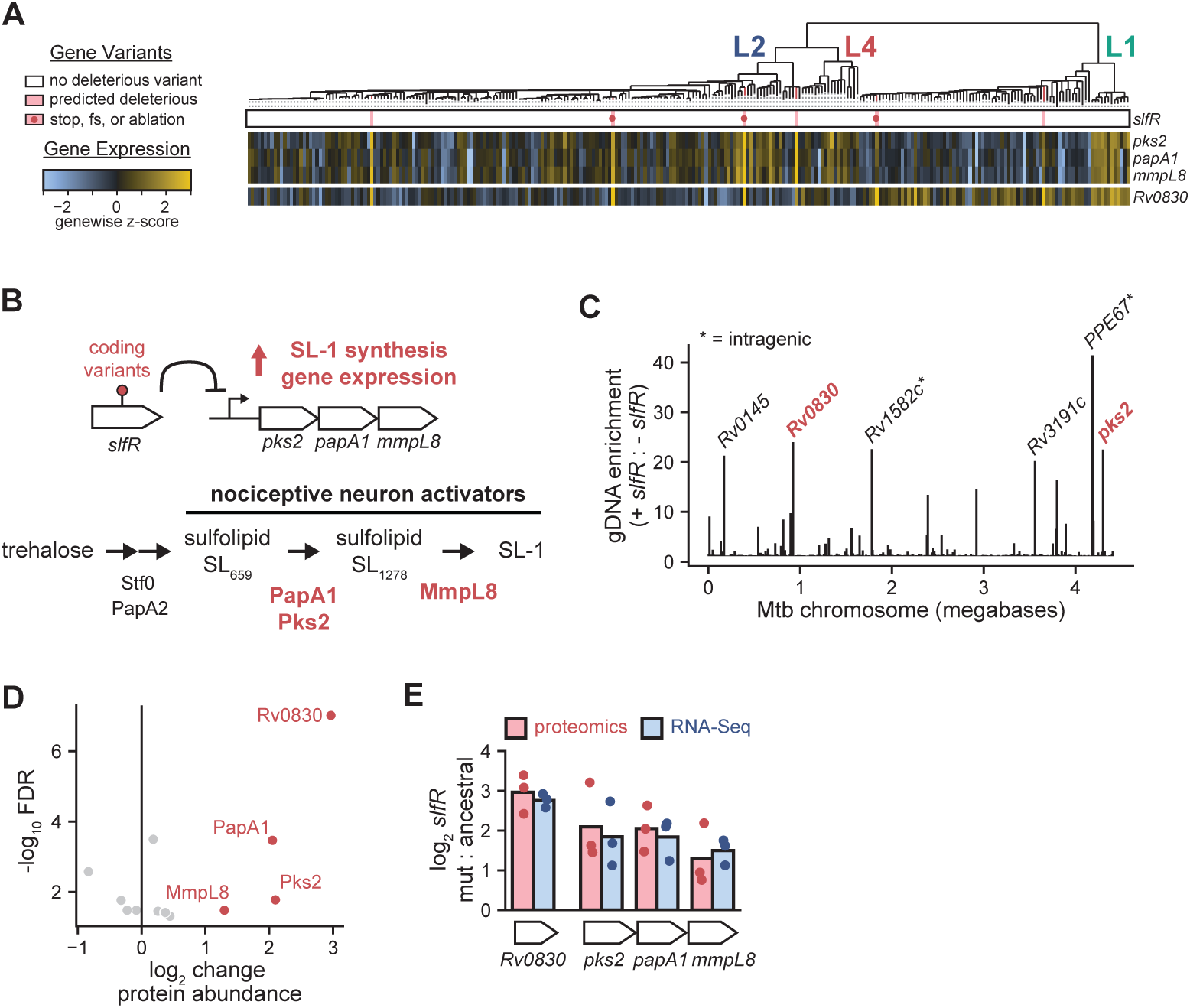
SlfR is a regulator of sulfolipid-1 biosynthesis gene expression. **(A)** Variants in *slfR* (*Rv0042c*) are plotted against a heatmap of expression z-scores of associated genes from GWAS. Red lines on the tree denote the inheritance of variants. Predicted deleterious, frameshift, and pre-stop variants are indicated. Strain gene expression is shown by heatmap relative to the gene’s mean across all strains; blocks of genes are co-operonic. **(B)** Schematics summarizing purposed SlfR regulation of *pks2-papA1-mmpL8* and showing role of the operon in biosynthesis of sulfolipid-1 (SL-1). **(C)** Bar plot showing mean enrichment from two replicates of Mtb gDNA incubated with or without purified SlfR. Genes nearby each peak are labelled. **(D)** Volcano plot showing z-test results for the mean of 2 genome-wide proteomics replicates for 3 pairs of closely related strains differing by *slfR* variants. Only genes passing an FDR of 0.05 are plotted and 4 genes with corresponding IDAP-seq peaks are labelled. **(E)** Bar plot summarizing change in protein and transcript abundance between three pairs of closely related isolates differing by *slfR* variants. Proteomics and transcriptomic datapoints shown are the average of replicates. See also Figure S4.

To test this predicted interaction, we purified SlfR and conducted *in vitro* DNA purification and sequencing (IDAP-Seq) to identify SlfR-bound regions of the Mtb genome^51^. The upstream intergenic region of the *pks2* operon was the 4^th^ highest binding peak in this assay, indicating direct transcriptional regulation (Figure 4C, Table S4). Comparison of the top binding regions with *meme* identified a ∼13 nucleotide palindromic SlfR binding motif (Figure S4A).

We further assessed this regulatory relationship by performing quantitative proteomics on three mutant-wildtype strain pairs that differed by a predicted deleterious *slfR* variant. The 4 greatest changes in protein abundance were Pks2, PapA1, MmpL8 and Rv0830, all of which were more abundant in the *slfR* variant strains (Figure 4D, Table S5). *Rv0830* encodes an uncharacterized methyltransferase not previously associated with sulfolipid production and was also identified as a direct target of SlfR by IDAPseq (Figure 4C). Comparison of transcript and protein abundance across the three selected strain pairs showed concordance across all 4 genes (Figure 4E). Together, these data show that in clinical strains, there is expression diversity in SL-1 synthesis and transport genes through the repeated acquisition of variants in *slfR*.

### Widespread *whiB6* variants associate with decreased secretion of ESX-1 virulence effectors

Finally, and most strikingly, the GWAS identified an association between decreased expression of *PE35* and *PPE68*, the first genes in the operon *PE35-PPE68-esxBA,* and recurrent predicted functional variants in the known ESX-1 transcriptional regulator, *whiB6*^52^. While *esxBA* were not captured in our stringent test for expression outliers, associations between decreased expression of *esxBA* and *whiB6* variants were confirmed by *pyseer* and inspection of the phylogenetic tree (Figure 5A, Table S6). Across our 274 strains, there were 13 independent *whiB6* variants which showed concomitant decreases in *PE35-PPE68-esxBA* operon expression (Figure 5B); 6 of these were inherited by one or more strains. We also identified 5 independent variants upstream of *whiB6* that coincided with a peak in mutational events in our global phylogenetic analysis (Figure S5A); a variant at this locus was previously described as altering *whiB6* function in a single strain^52^.

**Figure 5.**
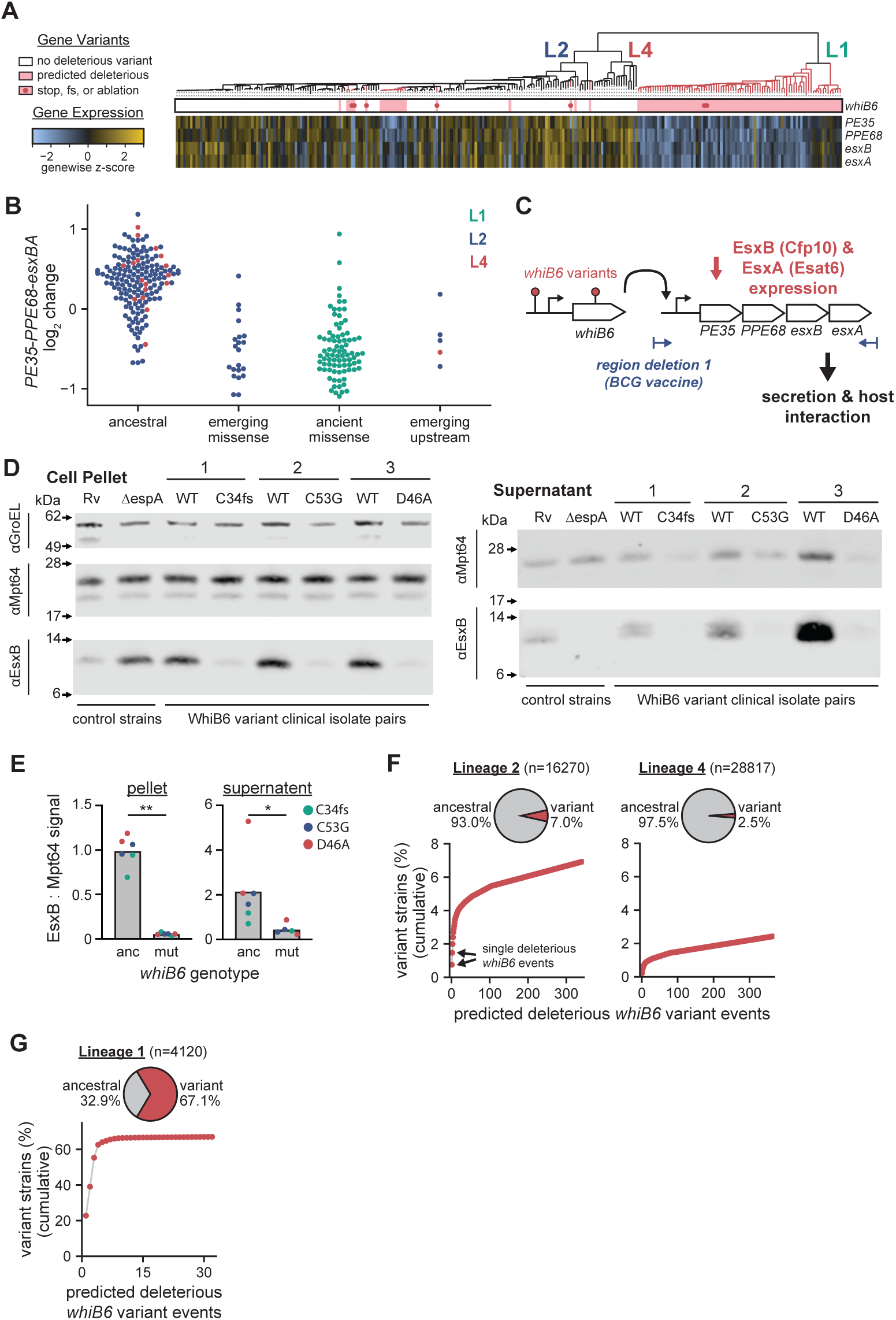
Frequent *whiB6* variants associate with decreased key ESX-1 virulence effectors. **(A)** *whiB6* variants plotted against a heatmap of expression z-scores of the *PE35-PPE68-esxBA* operon. Red lines on the tree denote the inheritance of variants. Plotted as in 4A. **(B)** Swarm plot showing replicate mean expression of *PE35-PPE68-esxBA* operon relative to the mean across all strains colored by lineage. Strains are categorized by *whiB6* genotype: emerging missense (predicted deleterious coding variants in lineages 2 and 4), ancient missense (1 of the 2 widely inherited predicted deleterious coding variants spanning lineage 1), emerging upstream (variants within 2 bases of the upstream peak in S5A), or ancestral (none of the prior variants). **(C)** Schematic summarizing purposed effect of *whiB6* variants on expression of ESX-1 effectors. **(D)** Western blots for EsxB (secreted effector), Mpt64 (secreted control), and GroEL (cytosolic control) across 3 closely related clinical isolate pairs differing by *whiB6* coding variants. H37Rv and H37Rv *ΔespA* are shown as controls to show loss of EsxB secretion without the EspA chaperone. Protein extracted from cell pellet and supernatant of Mtb grown *in vitro* are shown. **(E)** Quantification of EsxB and Mpt64 bands from 5D and S5B. Student’s t-test p-values=*0.016 and **<1e-4. **(F)** At the top, pie charts showing the strain frequency of predicted functional *whiB6* variant strains across lineages 2 and 4. At the bottom, cumulative distributions showing the cumulative fraction of predicted functional *whiB6* variant strains driven by each independent mutational event. **(G)** Pie chart and cumulative distribution of predicted functional *whiB6* variant strains plotted as in 5F but for lineage 1. See also Figure S5.

Recurrent evolution towards decreased expression of the *PE35-PPE68-esxBA* operon was unexpected: the genes *esxB* and *esxA* encode the key virulence effectors EsxB/Cfp10 and EsxA/Esat6, which are secreted as a heterodimer by the ESX-1 secretion system (Figure 5C)^53^. The importance of ESX-1 to Mtb virulence is well described. Loss of ESX-1 is the primary attenuating deletion in the vaccine strain, BCG^14^. The ESX-1 genes together and individually are required for wildtype growth of Mtb in animal models and macrophages, permeabilization and lysis of maturing phagosomes, and resulting induction of inflammatory cytokines including type I interferons and IL-1β^54–56^. EsxA and EsxB are also dominant T cell antigens, used as diagnostic targets in clinical IFNγ release assays, and are part of several of the vaccine constructs in late-stage development^36,37^. PPE68 has been shown to facilitate secretion of multiple ESX-1 substrates, including EsxA^57^.

To determine whether decreased *PE35-PPE68-esxBA* expression was sufficient to alter protein abundance, we selected 3 pairs of closely related strains differing by predicted deleterious *whiB6* variants and measured EsxB abundance in cell pellets and supernatants by Western blot. EsxB abundance in cell pellet was lower relative to control proteins in the pairwise comparisons of *whiB6* variant vs neighbor strains (Figure 5D, S5B). Abundance of secreted EsxB was also lower in *whiB6* variant strains. We note that secretion of Mpt64, a Sec-substrate, was also modestly lower in the variant strains. Nonetheless, the ratio of EsxB to Mpt64 was significantly lower in both the pellets and supernatants of the *whiB6* variant strains (Figure 5E), consistent with the model that *whiB6* variants result in decreased expression, abundance, and secretion of EsxB.

### ESX-1 regulatory variants arise frequently and are widely inherited in global isolates

To determine if *whiB6* variants are widespread globally, we returned to our phylogenetic analysis and ancestral reconstruction of 55,259 isolates. Across lineages 2 and 4, we observed >600 independent predicted functional *whiB6* mutational events accounting for 7.0% and 2.5% of strains, respectively; many of these variants subsequently expanded into multiple strains through clonal inheritance (Figure 5F). Strikingly, in lineage 1, 67.1% of strains have a *whiB6* variant driven primarily by 4 widely inherited alleles (Figure 5G); two of these lineage 1 ancestral alleles were present in our strain pool (Figure 5A). Across global isolates, the combination of recurrence of *whiB6* mutational events in lineages 2 and 4 and the prevalence of strains more ancient *whiB6* variants in lineage 1 results in widespread low ESX-1 virulence effector expression. These data challenge a naïve assumption that high virulence factor expression is uniformly beneficial to Mtb.

Mtb lineage 1 accounts for at least a quarter of TB cases worldwide but is less well studied than the other major lineages^58^. In lineage 1, we noticed a bimodal expression pattern of genes in the *espACD* operon, a *trans*-encoded part of the ESX-1 system, where the magnitude expression switched at two ancestral nodes on the tree (Figure 6A). The *espACD* operon is required for ESX-1 secretory activity and Mtb virulence in mice and macrophages (Figure 6B); EspA and EspC are also known Mtb antigens, and EspC is used in at least one vaccine candidate^59–61^. The *espACD* operon genes were also among our high expression diversity set. While our GWAS did not find coding variant associations with *espACD* expression, the GWAS did not assess non-coding regions. The *espA* upstream region is ∼1.2 kb and binds multiple regulators across its length including WhiB6, MprA, PhoP, and EspR^62–64^. Inspection *espACD* promoter region variants revealed two independent >1 kb upstream deletions matching the patterns of increased operon expression^65^. Mapping all *espA* upstream mutational events revealed that the sites of the deletions are also sites of frequent short variants at two sites (Figure 6C); these sites have a higher burden of genetic variation than >99% of intergenic windows across the genome (Figure 6D). One strain in our RNA-Seq library had a site B variant and showed higher *espACD* expression than outgroup strains (Figure 6E). Ancestral reconstruction of the upstream region of *espACD* revealed that most global lineage 1 isolates have either an upstream deletion or site B variant (Figure 6F). In contrast, only about 1% of lineages 2 and 4 strains have an *espACD* upstream variant with limited clonal inheritance (Figure S6A). Taken together, these data suggest that while ESX-1 components are under regulatory selection across Mtb strains, the pattern of inheritance and regulatory mechanisms under selection are distinct in each lineage, suggesting the impact of previously unrecognized genetic interactions.

**Figure 6.**
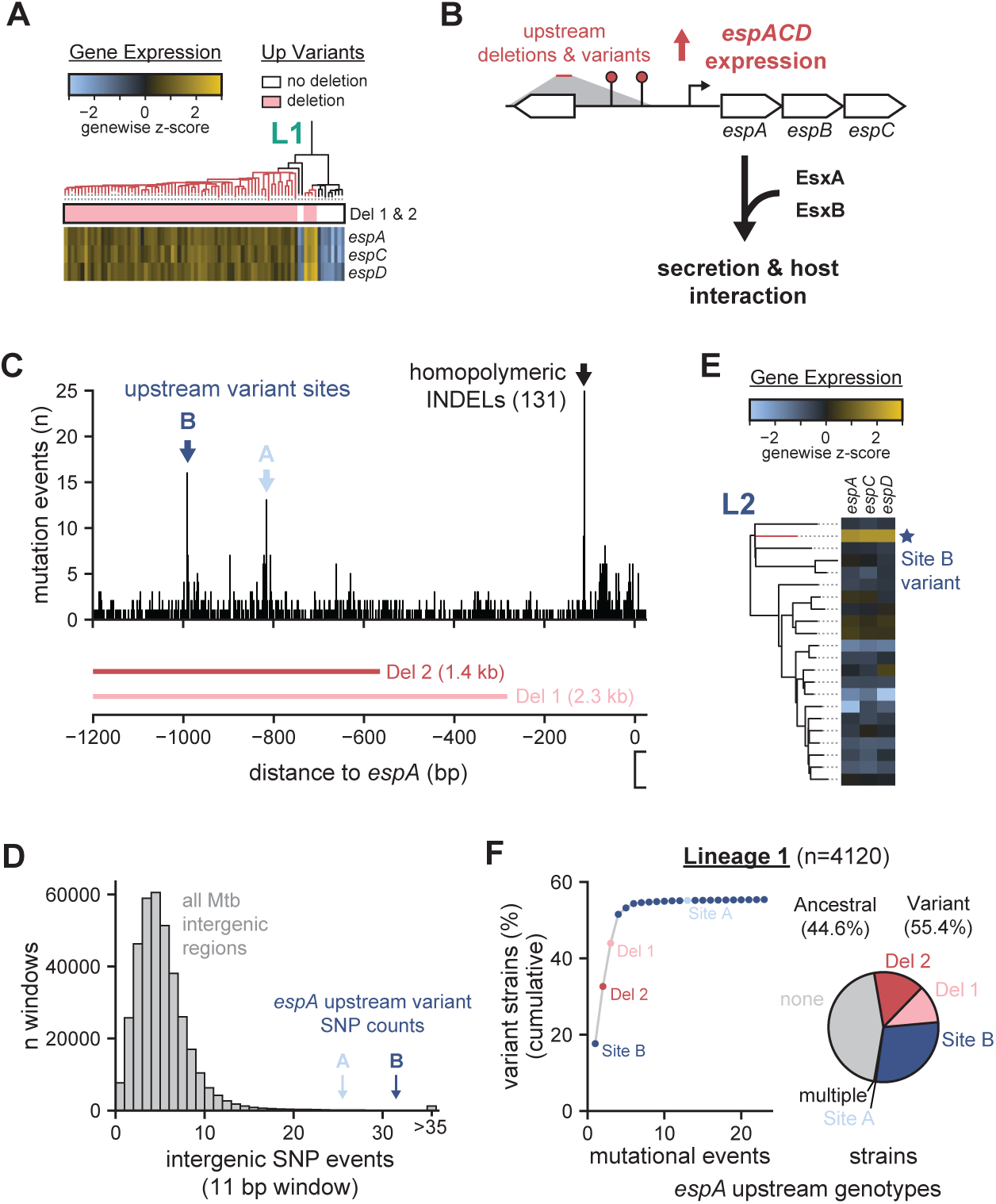
ESX-1 regulatory variation is widespread at the *espACD* locus in Mtb lineage 1. **(A)** Heatmap of *espACD* expression z-score in lineage 1 relative to the mean across all strains as in 4A. Variant row shows the presence or absence of deletions shown in 6C. **(B)** Schematic summarizing the purposed effect of *espACD* upstream variants on *espACD* expression. **(C)** Bar plot counting short (≤5 nt) mutational events across global isolates upstream of the *espACD* operon. Two upstream variant peaks are highlighted and as well as variation at a homopolymeric site. **(D)** Histogram showing the distribution of SNP events in all 11 base windows in intergenic regions with high quality variant calls. Position of *espACD* upstream peaks on the distribution are shown. **(E)** Heatmap of *espACD* expression for a clade of lineage 2 strains highlighting a single strain with a unique variant near Site B. **(F)** Pie chart and cumulative distribution showing inheritance of *espACD* upstream variants in lineage 1 plotted as in 5F. Slices and points are colored by variant shown in 6C. See also Figure S6.

### Frequent, modern *whiB6* variants associate with transmission of antibiotic resistance

One of the strongest evolutionary pressures facing Mtb is antibiotics. New cases of pulmonary tuberculosis are typically treated for 6 months with combinations of the first line drugs rifampicin, isoniazid, pyrazinamide, and ethambutol^66^. Variants that confer marginal or conditional survival advantages in drug facilitate survival and acquisition of high-level drug resistance, contributing to a pattern of recurrent emergence^67^. Therefore, we sought to assess the timing of *whiB6* and *slfR* coding and *espA* promoter variants relative to the introduction of antibiotic pressure. To measure the relative modernity of our variants, we compared the distance to root of variants to the null distribution of randomly chosen branches weighted by length (Figure 7A, Table S7)^68^. To account for lineage differences in phylogenetic structure, we separated our analyses by lineage^69^. In lineages 2 and 4, *whiB6* variants are modern relative to the null distribution of variants and in lineage 4 *slfR* variants are modern. However, in lineage 1, both *whiB6* coding and *espA* upstream variants are more ancient than the null distribution, suggesting that antibiotic alone does not account for the emergence of these variants.

**Figure 7.**
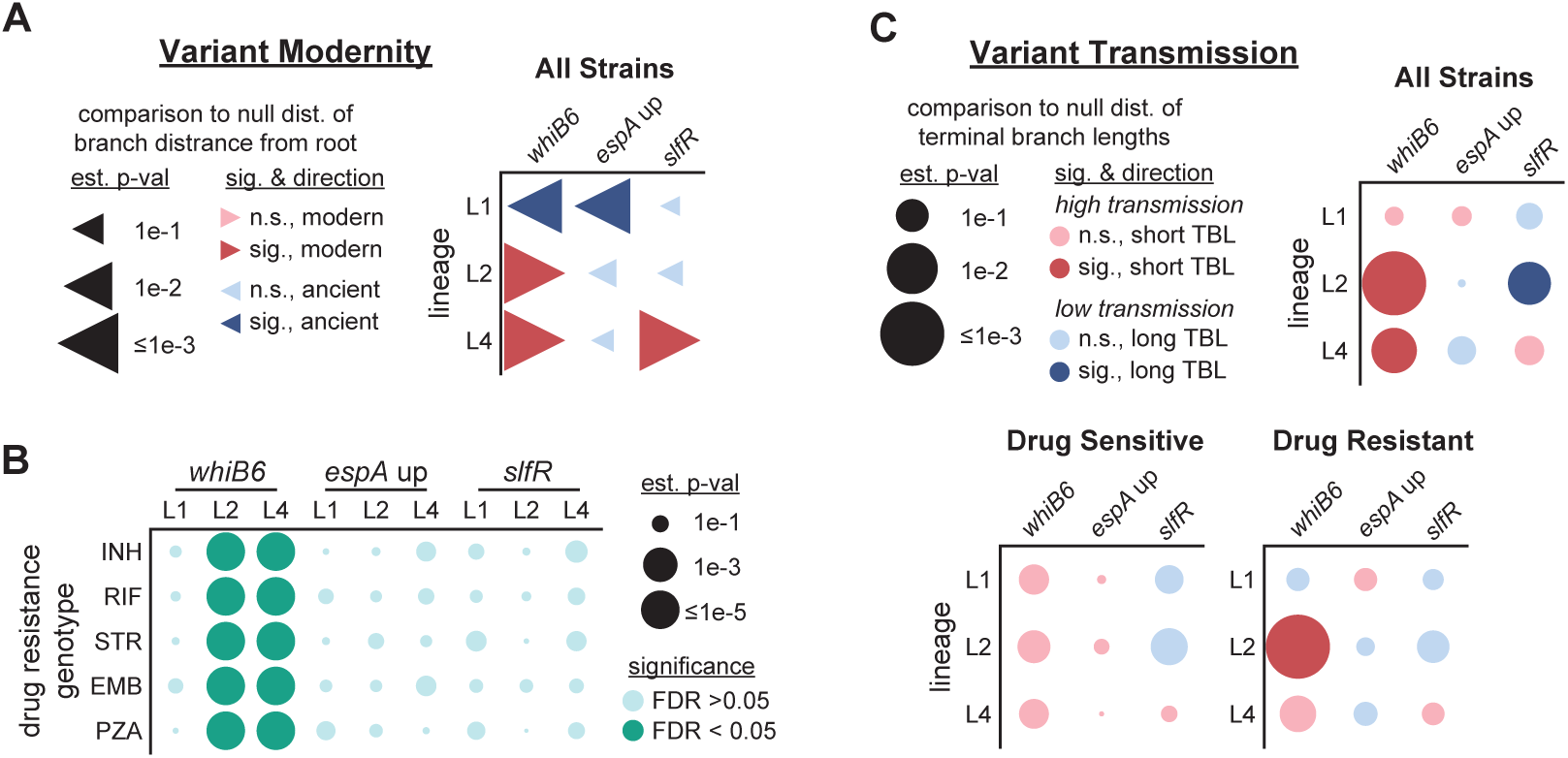
Modern *whiB6* variants associate with drug resistance and transmission. **(A)** Statistical association of predicted deleterious *whiB6* and *slfR* variants and *espACD* upstream variants with modernity, measured by distance from root and compared to a null distribution of randomly selected branches. Color denotes the directionality and significance (estimated p-value < 0.05) of associations. Size indicates significance of association. **(B)** Statistical association of variants from 7A with selected high level drug resistance across global isolates separated by lineage. Dot size shows estimated p-value and color denotes significance with Benjamini-Hochberg FDR α=0.05. **(C)** Comparison of mean terminal branch length, a metric of transmission, of clades with variants from 7A to a null distribution calculated by random permutations. Associations were run on the complete tree or with trees with strains with genotypic drug resistance or drug sensitivity. Color denotes the directionality and significance (estimated p-value < 0.05) of associations. Size indicates significance of association.

To more directly test the relationship of our variants with drug pressure, we quantified population genetic links between our variants and drug resistance genotypes. Mirroring the temporal analysis, we found strong associations between *whiB6* variants in lineage 2 and 4 and known drug resistance mutations (Figure 7B, Table S7). Critically, because our transcriptomic cohort only included phenotypically drug sensitive isolates, we know that *whiB6* variants alone do not cause high-level drug resistance *in vitro*. These data suggest that in lineages 2 and 4, *whiB6* variants may either contribute to acquisition of drug resistance or increase the fitness of drug-resistant strains.

Transmissibility is critical for strain success. Therefore, we assessed the relationship between our variants and transmission. We measured the mean terminal branch length, a metric of transmission, of clades or strains where variants evolved and compared to random permutations (Figure 7C, top; Table S7)^11,70,71^. Matching our drug resistance associations, *whiB6* variant clades have significantly shorter terminal branch lengths in lineages 2 and 4, pointing towards increased transmission. To determine if this effect is specific to drug resistant strains, we ran the association separately on trees with only or without genotypic drug resistance (Figure 7C, bottom). In this analysis, *whiB6* variants clades maintained significantly shorter terminal branch lengths in drug resistant lineage 2 (est. p-value <1e-5) and trended towards shorter lengths in drug resistant lineage 4 and drug sensitive lineage 2 and 4 strains (est. p-values 0.068, 0.10, 0.14). Taken together, our data links *whiB6* variants to increased transmission of drug-resistant strains in some Mtb genetic backgrounds, but suggest that *whiB6*, *slfR*, and *espA* promoter variants can also be driven by pressures other than drug.

## Discussion

### Expression diversity reveals paradoxical selection on ESX-1 secreted protein expression

The formula for evolutionary success in the age of antibiotic selection is different than in the pre-antibiotic era. Drug is a modern pressure, and multiple studies have found that Mtb is evolving many forms of altered drug susceptibility beyond high-level drug resistance^5,44,68,72^. We identified an association between variants in the ESX-1 transcriptional regulator, *whiB6*, high-level drug resistance and the ongoing transmission of resistant strains (Figure 7). Our data suggests *whiB6* variants provide a selective advantage during drug treatment and thus increase the rate of acquisition of resistance variants^67^.

Unexpectedly, these recurrent *whiB6* variants *downregulate* essential ESX-1-secreted virulence factors in the *PE35-PPE68-esxBA* operon (Figure 5). The importance of the ESX-1 locus for Mtb virulence is clear in macrophages, in mice, and—most importantly—in people where loss of ESX-1 is the mechanism of attenuation of the BCG vaccine strain. EsxA and EsxB have well defined roles in the permeabilization of the macrophage phagosome and induction of type I interferons, which are associated with increased bacterial burden and severe disease^14,54–56,73,74^. A tempting model is that *whiB6* variants arise due to a selective pressure to decrease virulence; for example, by inducing less severe or asymptomatic disease, variant strains may evade treatment through avoiding TB surveillance efforts, thus providing a modern selective advantage and facilitating spread. Alternatively, the source of selective pressure may be altered expression of another WhiB6 target whose selective advantage outweighs downregulation of *PE35-PPE68-esxBA*^75^.

We also found expression variation in ESX-1 secreted proteins has occurred via distinct evolutionary paths in various Mtb lineages (Figure 5G, 6F). Driven by ancestral variation, a majority of lineage 1 strains harbor variants we associate with decreased *PE35-PPE68-esxBA* expression or increased expression of the *espACD* operon, which encodes additional ESX-1-associated proteins. Though historically understudied due to limited geographic distribution and lower virulence, lineage 1 accounts for at least a quarter of TB cases worldwide^4,58^. Recently, it has been associated with longer infective periods and, combined with lower virulence, suggested to play a role in asymptomatic TB^4,76^. Future work will be required to determine if lineage 1’s prevalent *whiB6* and ESX-1 variation contributes to these phenotypes. Regardless, the differential evolutionary pressure and timing of ESX-1 variation in lineage 1 vs lineages 2 and 4 shows there are different formulas for Mtb’s success.

### Expression diversity as a mechanism to modulate host interaction

Expression variation in ESX-1 is part of a wider trend of variation across Mtb–host interactive processes. The recurrent emergence of expression variation in *in vivo* essential genes suggests natural selection (Figure 3A). Coupled with enrichment of expression diversity in known antigens, we propose that expression diversity is a mechanism for Mtb to tune function of genes at the host-pathogen interface without altering their coding sequences^6^.

In *slfR* variants, we identified a selective pressure towards increasing expression of SL-1 synthesis genes; in lineage 4, where *slfR* selective pressure was strongest (pN/pS = 1.95), these variants were primarily modern but without a clear link to drug selection (Table S1, Figure 7). Early work identified sulfolipid abundance as highly variable between isolates and positively correlated with lung bacterial burden^49^. With SL-1’s recently elucidated role as an activator of nociceptive neurons to induce cough, our data supports a model in which *slfR* variants modulate host interaction^50^. We also identified high expression diversity in 2,3-diacyltrehaloses (DAT) and penta-acyltrehaloses (PAT) synthesis genes (*pks3-pks4-papA3-mmpL10;* Figure S3A), highlighting regulation of immunomodulatory lipid synthesis genes as a potentially rich source of isolate phenotypic variation^77^.

These data have critical implications for TB interventions. Though the genetic essentiality of Mtb genes to pathogenesis has been considered, regulatory diversity—in particular, tolerance of decreased expression of essential virulence genes—has not been explored. These concerns are not theoretical: the safety of the MTBVAC vaccine, now in clinical trials, is supposed to be achieved through deletion of the *phoPR* regulator, resulting in reduced ESX-1 effector expression^78^. ESX-1 effectors we identified as expression diverse are key antigens involved in TB diagnostics and targets for vaccine design; 6/14 (43%) of novel vaccine antigens were among our expression diverse set. Inherited and *de novo* variation in antigen abundance across different strains could cause variation in the efficacy of vaccines and the specificity of TB tests relying on immune response to whole Mtb (PPD skin test) or specifically EsxB/EsxA (IGRA blood test)—indeed, a few strains with these phenotypes have been described but interpreted as the exception rather the rule^79,80^.

### Towards characterizing a diverse and evolving transcriptome

To analyze the evolution of Mtb gene expression, we made several technical advances. First, we built a cost effective, scalable RNA-Seq technique for bacterial samples or any non-poly-adenylated RNA. A core innovation in our method is the addition of a programmable ssDNA depletion technique using DNA oligos and Taq polymerase’s endonuclease activity; depletion acts on pooled and barcoded cDNA, removing the need to conduct rRNA depletion on individual samples or unstable barcoded RNA (Figure 1A, S1C). Through this, we reduced the cost of library preparation to ∼$5 per sample and eliminated single-sample handling from the bulk of the protocol. In addition to linking genetic variants to expression diversity, bringing phylogenetic-transcriptomic analyses to this scale enabled us to identify transcriptional divergence across long and short evolutionary time scales (Figures 2B, 2C). Prior studies have focused on exploring diversity across distant strains and lineage boundaries, but our data highlight that major transcriptional differences can arise in recently diverged strains and clades^9,10^. An extreme example of a burst of expression variation is in the recently evolved (circa 1950), highly transmissible Peruvian clade visually separable by expression heatmap and PCA (Figures 2B far left)^12^. Application of machine learning models to this scale of transcriptomic data will facilitate identification of transcriptional signals that are specific to successful strains and clades^81^.

To better understand Mtb’s global phylogenetic diversity, we constructed a parallelizable pipeline for genome wide variant calling, quality control, tree building, and ancestral reconstruction on tens of thousands of Mtb clinical isolate sequences. In addition to supporting our conclusions around regulatory selection, this pipeline produced phylogenomically informative data for all Mtb genes. These data include (1) gene pN/pS values to measure the directionality of selection (Table S1), (2) summaries of the acquisition events and inheritance of all identified variants, and (3) the global phylogenetic tree with variant event nodes, all of which are available as community resources.

These data demonstrate that Mtb is not a genetically or phenotypically monomorphic organism. Modern advances in high throughput phenotyping, genomics, and transcriptomics give us the opportunity to analyze strains at scale and future study of the TB global pandemic must encompass both existing and evolving diversity and be aware of the limitations of laboratory strains.

### Limitations of this study

Transcriptional pathways are responsive and can be activated by host stressors such as acid, hypoxia, or nutrient conditions^81^; our study, conducted under *in vitro* growth, cannot be viewed as a complete picture of the possible regulatory change across Mtb isolates. We cannot rule out the possibility that transcriptional effects on *PE35-PPE68-esxBA*, *espACD*, and *pks2-papA1-mmpL8* may be further modulated *in vivo*. Future work will be required to mechanistically untangle the effects of regulatory evolution *in vivo*. Regardless, the repeated evolution and population success of strains with our regulatory variants is clear.

This study relied on GWAS tools to associate transcriptional variation in Mtb clinical isolates with genomic variants; associations do not always indicate causality. Furthermore, we made use of predictive software to classify coding variants as functional^82,83^. We and others have observed variation in evolution pressure across the Mtb phylogenetic tree (Table S1)^6^; suggesting introduction of genomic variants into laboratory strains may remove important epistatic interactions present in the isolate lineages in which they evolve. Within this study, we addressed this limitation by comparing multiple sets of closely related isolates whose only shared difference was the variant of interest (Figures 4D, 5D); though outside of the scope of this study, future work should expand Mtb genetic tools to work in arbitrary clinical isolates.

This study uses public short read sequencing data. Though this scale provides critical statistical power, the provenance of this data can vary—for example, isolates could be from active or passive cohorts or be enriched for drug resistant strains. In this work, we removed studies with clear undesirable biases (e.g. resampling patients over treatment). Our transmission analyses used terminal branch lengths, an established metric in the Mtb field, and explored drug effects by separating resistant and sensitive trees (Figure 7)^11,70,71^. To limit distortions driven by differences across the phylogenetic tree, we separated most analyses by bacterial lineage^69^. At a minimum, our global isolate data strongly supports our assertion that hypomorphic ESX-1 variants (1) are widespread, (2) occur repeatedly, and (3) are readily transmitted in clinical isolates, all of which are unexpected for variants in this canonical virulence pathway.

## Supporting information

TableS1 pNpS

TableS2 expression diversity

TableS3 variant associations

TableS4 SlfR binding peaks

TableS5 SlfR proteomics

TableS6 esxBA expression associations

TableS7 phylogenomics analyses

TableS8 WGS accessions

TableS9 primers and reagents

## Resource Availability

De-multiplexed MTB RNA-Seq and IDAP-Seq datasets are uploaded to GEO (GSE294801, GSE294802). Data processing and plotting was conducted in Jupyter notebooks; code used to produce data figures and links to custom scripts are available (github.com/peterculviner/HT-Mtb-RNA-Seq-Paper-Notebooks). Large datasets including trees, variant calls, ancestral reconstruction calls, and processed MTB RNA-Seq and IDAP-Seq counts are uploaded to Mendeley (DOI: 10.17632/h9dd6hsvc2.1). Whole genome sequencing for the transcriptomic strains were previously uploaded under PRJNA1039243 (Vietnam) and PRJNA1039243 (Peru). Individual whole genome sequencing accession numbers are in Table S8. Primers and reagents used in the study are in Table S9.

## Acknowledgements

This work was supported by a fellowship awarded to P.H.C. (1F32AI174653); a fellowship awarded to A.M.F. (DGE 2140743); and P01AI143575, R01AI184469, and R21AI187884 awarded to S.M.F. We thank Michael Laub for a critical reading of the manuscript.

## Author Contributions

P.H.C and S.M.F designed the study. P.H.C., A.M.F., and S.M.F. designed experiments. P.H.C. and A.M.F. conducted experiments. P.H.C., A.M.F., and Q.L. conducted analyses. P.H.C. and S.M.F. wrote the paper. D.T.M.H., P.V.K.T., D.D.A.T., N.L.Q., R.C., L.L., M.C., S.J.D., M.B.M., and N.T.T. provided isolate samples and metadata.

---

**Supplementary Figure 1.**
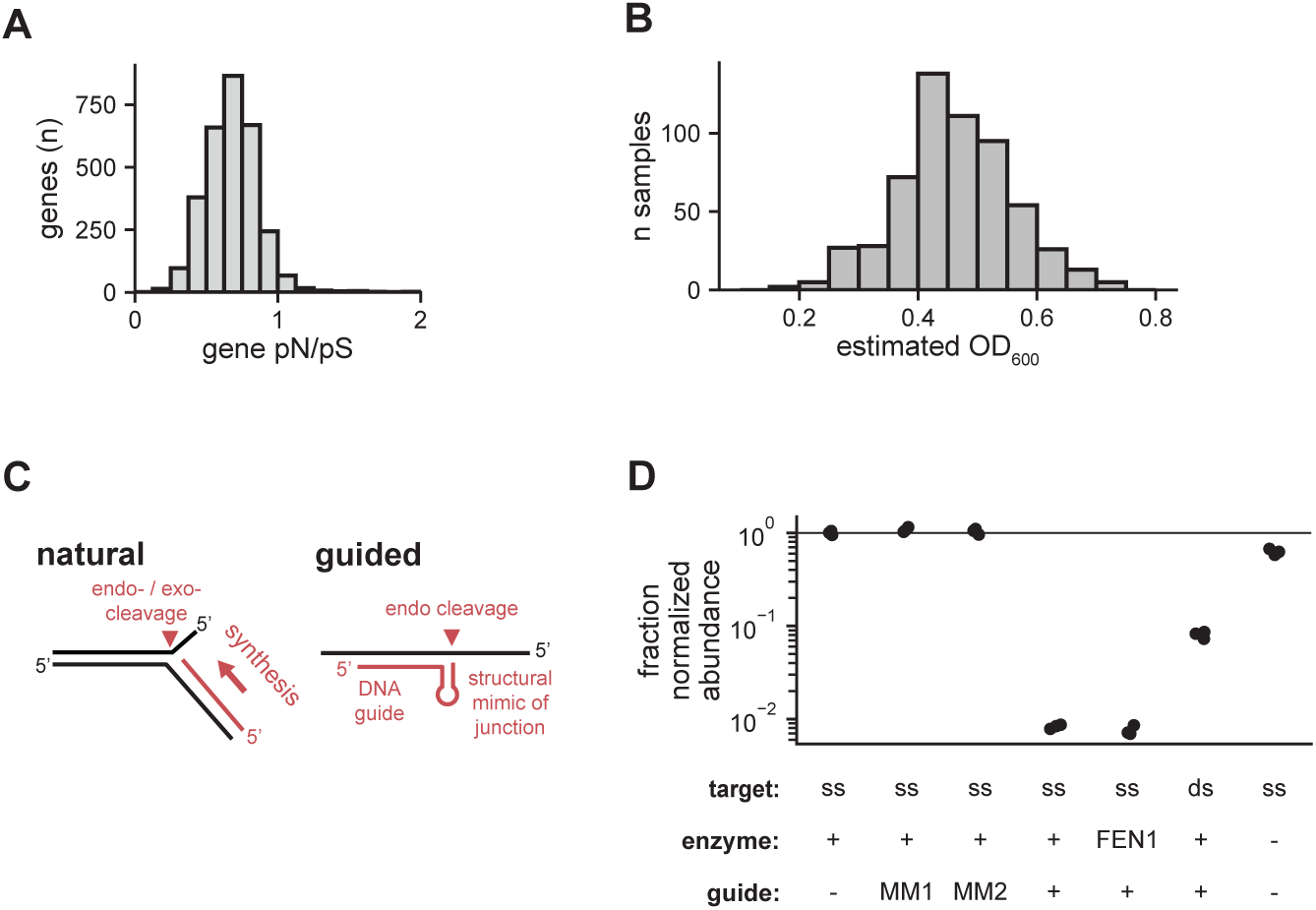
Multiplex analyses of Mtb clinical isolates and development of transcriptomic pipeline. **(A)** Histogram of pN/pS values for the lineage 1-4 phylogenetic tree calculated for all Mtb genes with 95% high quality calls and at least 5 synonymous and non-synonymous variants. **(B)** A histogram of the estimated optical density of all samples at the time of RNA harvest. **(C)** Schematic of Taq polymerase endonuclease activity during natural DNA synthesis and guided cleavage. **(D)** Log-transformed no guide control normalized qPCR of a single-stranded test oligo or mismatched controls (MM1/2), using a FEN1 endonuclease instead of Taq polymerase, with the addition of a complementary test oligo (ds), and a no enzyme control.

**Supplementary Figure 2.**
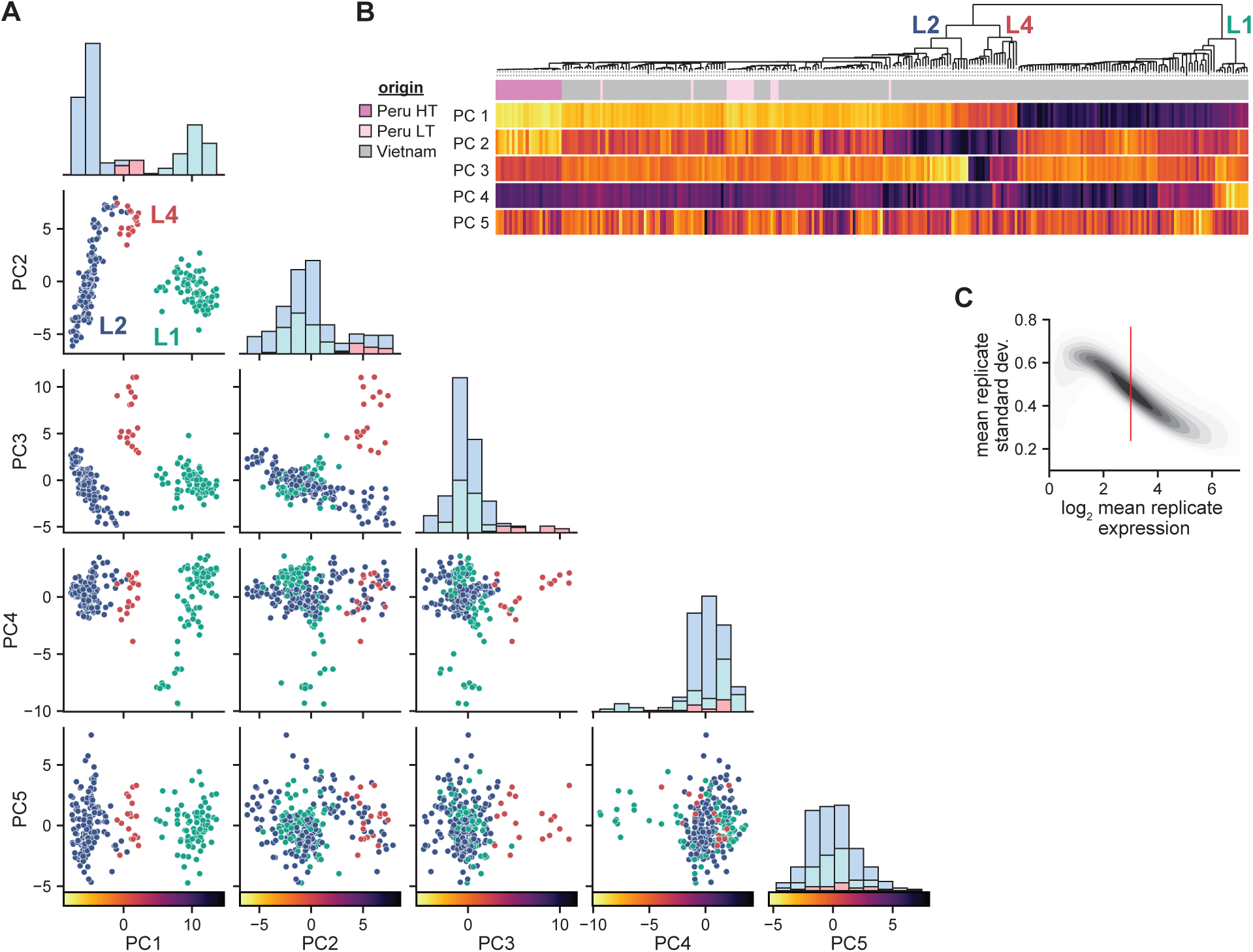
Deeper principal components separate Mtb sublineages. **(A)** Scatterplots of principal component analysis are shown as in 2C with all pairwise relationships for PC1-5 shown. Histograms of each component are also shown on the diagonal. On the x-axis label, colormaps legends are shown for heatmaps of PC1-5 against the phylogenetic tree in S2B. **(B)** Phylogenetic tree of Mtb isolates with transcriptional data with a heatmap of strain PC 1-5 values from S2A plotted against all strains as well as source cohort. **(C)** Kernel density plot of the mean standard deviation between replicates vs the mean expression level. Used to pick an expression threshold that minimizes replicate variation; threshold of 3 shown in red, includes 54% of coding regions.

**Supplementary Figure 3.**
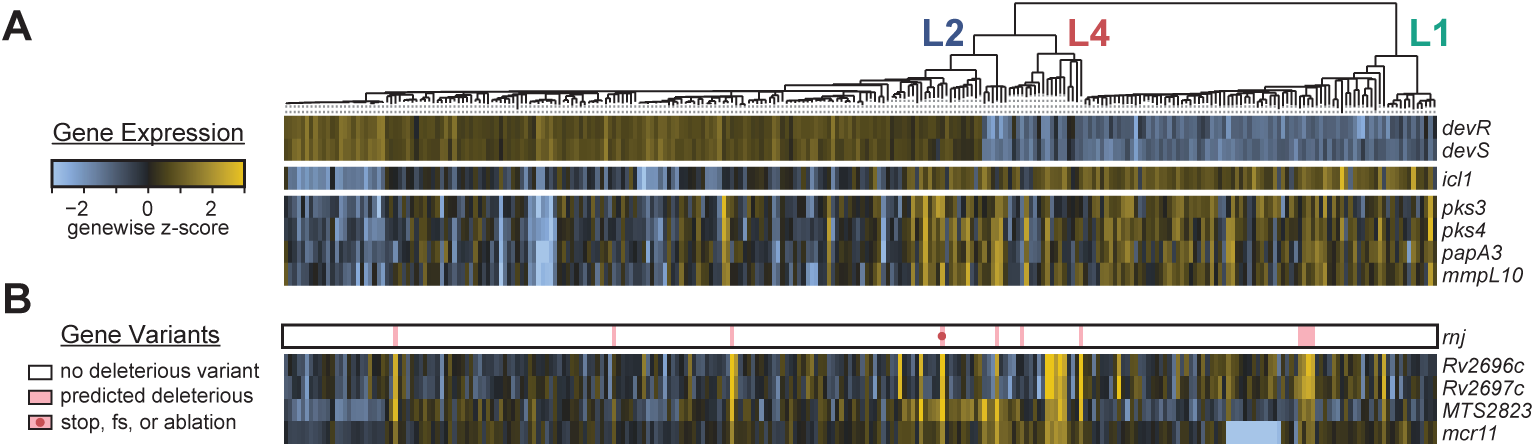
Additional genes with diverse expression across Mtb isolates. **(A)** Phylogenetic tree of Mtb isolates with a heatmap of expression z-scores from selected genes discussed in the text. Strain gene expression is shown by heatmap relative to the gene’s mean across all strains; blocks of genes are co-operonic. **(B)** Variants in *rnj* plotted against a heatmap of expression z-scores of associated genes from GWAS. Predicted deleterious variants are shown in pink with frameshift and pre-stop variants indicated by a circle.

**Supplementary Figure 4.**
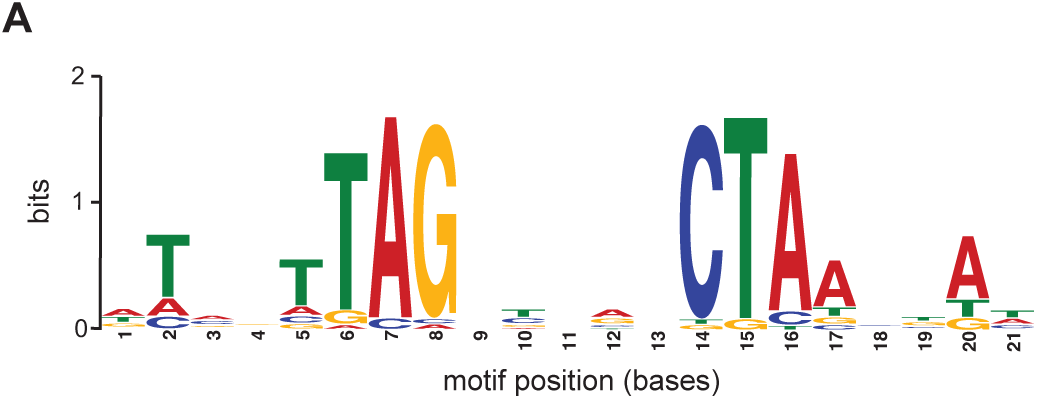
SlfR binding motif. **(A)** SlfR binding motif identified by *meme* analysis of top IDAP-Seq peak regions.

**Supplementary Figure 5.**
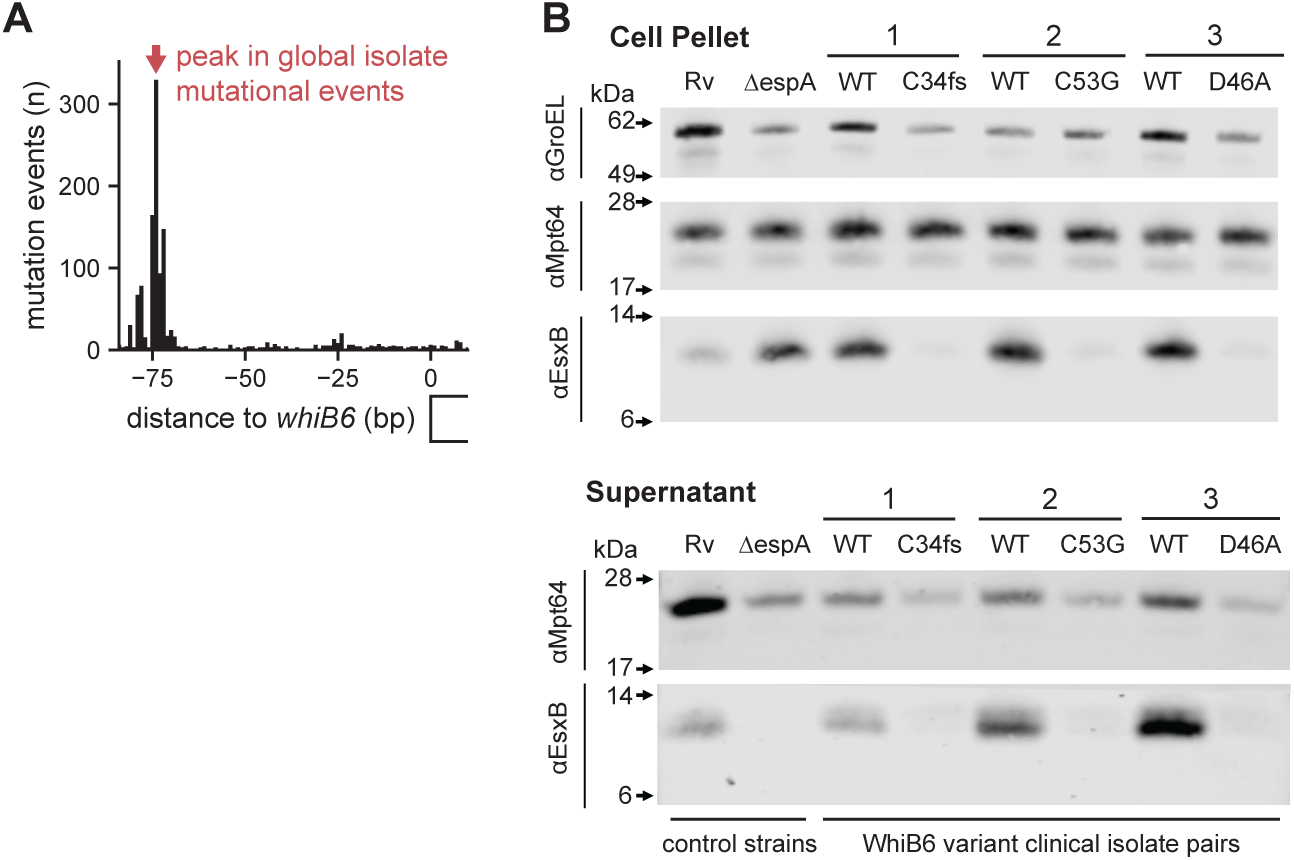
Replicate blots and *whiB6* upstream variants. **(A)** Bar plot counting mutational events across global isolates upstream of the *whiB6* gene. **(B)** Biological replicate of 5D.

**Supplementary Figure 6.**
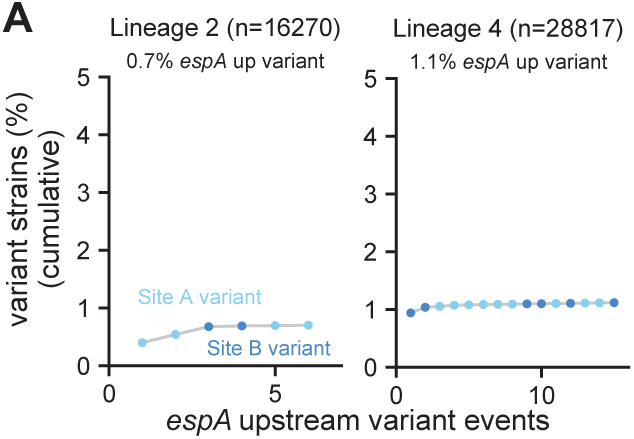
e*s*pACD upstream variants in lineages 2 and 4. **(A)** Cumulative distributions showing inheritance of *espACD* upstream variants in lineage 2 and 4 as in 6F.

## Supplemental Table Titles

**Table S1. pNpS by lineage.**

**Table S2. Gene expression diversity across Mtb lineages.**

**Table S3. GWAS of expression diverse strains.**

**Table S4. Top SlfR binding peaks by IDAP-Seq.**

**Table S5. Proteomics with *slfR* variant strains and nearest neighbors.**

**Table S6. Association of genomic variants with *esxBA* expression.**

**Table S7. Association of *whiB6*, *espACD* upstream, and *slfR* variants with modernity, drug resistance, and transmission.**

**Table S8. Accession numbers of clinical isolates WGS used in this study.**

**Table S9. Primers and reagents.**

## Methods

### Clinical isolate data processing and variant calling

To identify samples to run through our pipeline we used *entrez* to identify the set of unique biosamples from 603 bioprojects with Mtb WGS sequencing data. We used *sra-tools* to download and convert data from the short read archive (SRA) to the FASTQ format; biosamples with multiple runs were concatenated and variant calling was run on the Harvard FAS Research Computing (FASRC) cluster.

Our variant calling pipeline did the following steps for each biosample. First, it downloaded the dataset from the short read archive using *sra-tools* and randomly down sampled large FASTQs to a target depth of 100x. Next, it used *fastp* to infer and remove adapters and duplicate reads. Next, *bwa mem* was used to map reads to the H37Rv (NC_000962.3) genome. Then, *tbprofiler* was used to call Mtb lineage-defining SNPs. Finally, fixed variants were called with *breseq* and mixed variants were called with *mutect2*. While the pipeline progressed, samples were removed (early stop) if any of the following was identified: the sample had an average depth less than 25, coverage of less than 80% of the Mtb genome, or *tbprofiler* did not call the strain as a member of Mtb lineages 1-9.

Next, we concatenated quality control metrics from all successful pipeline runs (n=62639) and removed low quality or possible “mixed infection” samples based on the following filter metrics: (1) greater than 3 standard deviations from the mean number of low quality sites called by *breseq*, (2) greater than 3 standard deviations from the mean error rate called by *samtools*, (3) less than 98% genome coverage, (4) *mutect2* called a high proportion of “mixed” single nucleotide variants (less than 75% of single nucleotide variants called by *mutect2* had an allele frequency greater than 90%), or (5) had greater than 2 mixed lineage defining variants according to *tbprofiler*. After filtration, 89.9% (n=56310) of input biosamples remained.

### Clinical isolate phylogenetic tree construction

For computational feasibility, the global isolate tree was constructed with a divide-and-conquer approach using *fasttree2*. Strains called by *tbprofiler* as in the sub-lineages 1.1, 1.2, 2.1, 2.2, 3, or 4.1-9 were grouped and used to build 14 separate sub-lineage trees; a scaffold set including a selection of strains from each sub-lineage was also collected (n=514). A *Mycobacterium canettii* outgroup was included in each set for rooting. Each of the sub-lineage sets and the scaffold set were run though our genome-wide SNP alignment pipeline to generate a fasta file for tree building. Briefly, *breseq* variant calls (or a missing data character) were written for all genomic sites with a SNP in at least one sample excluding (1) sites with missing data for more than 50% of samples or (2) sites that fell into poor variant recall regions.

All 15 trees were then constructed using *fasttree2* (compiled with -DUSE_DOUBLE) using the *rawdist* flag suggested for SNP alignments. To make tree branch lengths comparable, branch lengths were converted from substitutions per site (SNP sites) to substitutions per genome (all Mtb genome sites). Finally, the subtrees were converted into a global tree of lineages 1-4 by pruning and replacing a given subtree’s down sampled clade on the scaffold tree with the complete lineage subtree.

The transcriptomic cohort was small enough (n=274), that maximum-likelihood tree building was possible on all strains. The genome-wide SNP alignment pipeline was used as above to generate a fasta alignment for all strains and *Mycobacterium canettii* except for a more stringent missing data threshold of 10%. A generalized time-reversible (GTR) consensus tree was generated with *iqtree*, 1000 ultrafast bootstraps, and invariant genomic site counts.

### Clinical isolate ancestral reconstruction

We ran parsimony ancestral reconstruction separately on global isolates and the transcriptomic cohort using a custom pipeline. For each cohort, all isolate *breseq* variant calls and missing data were merged into a single variant call file (VCF) with a single variant per row using *bcftools*. For each variant, annotations and effects were called with *snpeff* and *sift4g*. Variant calls were converted into a custom format, variant call bytestream (VCB), enabling random access by either variant position or sample. Using VCB files and our phylogenetic trees, we conducted parsimony ancestral reconstruction separately for all variant rows with *pastml* and wrote a new VCB file describing the ancestral state at every node. Finally, nodes with unambiguous evolutionary events (Ref → Alt or Alt → Ref) and reversions were called using the ancestral VCB and phylogenetic tree and custom code to generate a final event VCB.

### pN/pS ratio calculation

Transition-transversion rate was estimated at 1.95 using unambiguous parsimony evolutionary events on the global phylogenetic tree. pN/pS ratios for each gene were calculated using the ratio of events for each coding region and calculated separately for the whole tree or each separate lineage: (observed nonsynonymous / possible nonsynonymous) / (observed synonymous / possible synonymous). Possible counts were calculated with *biopython* using the NG86 method^84^. Codons that overlapped with sites with (1) a missing data rate above 5% or (2) poor variant recall regions were excluded from the analysis—the included fraction of a CDS is recorded in Table S1.

Gene category enrichments in Figure 1B were calculated on the set of CDS with (1) an included fraction of 1.0 and (2) at least 5 observed nonsynonymous and synonymous events in all 4 major lineages (n=2803). Two-sided enrichment was calculated against each *mycobrowser* category and previously identified transcription factors using the Fisher exact test for each category. Previously identified transcription factors were defined as the union set of predicted transcription factors from prior studies of the Mtb regulatory network^25^.

### Mtb growth and RNA harvest

Mtb was grown in a 1 mL volume in 24-well plates, covered with breathe-easy film, at 37°C x 100 RPM, in 7H9 media supplemented with 0.2% glycerol, 0.05% Tween-80, and 10% OADC. Each Mtb strain had different growth and lag phase kinetics. To capture all strains at a similar OD_600_ (∼0.4), we separated growth into three phases: recovery, outgrowth, and harvest. OD_600_ values were estimated by well opacity measured by photography and comparison to photographs of a known OD_600_ standard curve. For recovery, 50 uL of thawed culture was inoculated into 950 uL of fresh media and grown for 72 hours. For outgrowth, OD_600_ was measured, and strains were back diluted to 0.03 in fresh media or allowed to continue growth if OD_600_ was less than 0.05; strains were allowed to grow for an additional 72 hours. For harvest, strains were back diluted to 0.03 to 0.1 depending on strain growth during outgrowth and allowed to grow for a final 48 hours. To halt transcriptome dynamics and sterilize cultures, 145 uL of 16% PFA was added to each 1 mL culture and allowed to incubate for 1 hour at room temperature before removal from the biosafety level 3 facility.

In 1.5 mL tubes: fixed cells were centrifuged for 10,000 RPM for 5 minutes, washed into PFA quencher (25 mM Tris HCl pH 7.5, 50 mM NaCl, 250 mM glycine, 0.1% Tween-20), pelleted again, and resuspended in 20 uL T2E (10 mM Tris pH 8, 2 mM EDTA). Crosslinks were reversed by incubating for 30 minutes at 70°C in strip tubes. Cells were lysed by adding 600 uL Trizol, transferring to a bead beating tube and bead beating 2 times at 6,000 RPM for 30 seconds. To separate Trizol, 120 uL of chloroform was added to each tube and vortexed. Tubes were centrifuged for 10 minutes at 10,000 RPM and the aqueous upper layer (250 uL) was extracted to a deep well 96-well plate. To precipitate RNA onto beads, 5 uL of undiluted carboxylate-modified magnetic beads and 260 uL of isopropanol-GuHCl (1 M guanidium-HCl, 0.05% Tween-20, in 100% isopropanol) was added and mixed with a multichannel. After a 5-minute incubation, plate was placed on a magnetic rack and allowed to clarify. With a multichannel, beads were pelleted and washed 4 times with 500 uL 90% ethanol, returning to magnetic rack after each wash. After the last wash, beads were allowed to dry for 5 minutes and resuspended in 15 uL T.1E (10 mM Tris 8, 0.1 mM EDTA). After incubation, RNA solution was placed in fresh 96-well plates for storage at −80°C.

### qPCR validation of Taq depletion

Feasibility of Taq depletion was tested in 30 uL reactions with or without 1 uL Taq or thermostable FEN1 with appropriate buffer, 2 uL 25 nM target primer (ss or ds), with or without 2 uL 250 nM guide, alternative guide, or mismatch guide. Reactions were incubated at 95°C for 30 seconds followed by 65°C for 1 hour. To degrade enzymes, 1 uL of thermolabile proteinase K was added and incubated 37°C for 15 minutes and inactivated at 60°C for 15 minutes. Triplicate reactions were back diluted 1:100 and 2.5 uL was added to 10 uL triplicate qPCR reactions using iTaq master mix and 300 nM primers. Relative quantity compared to no guide control was calculated using C_t_ of standard curve of target oligo dilutions.

### Design of Taq depletion primers

Primers targeting the rRNA (23S, 16S, 5S), tmRNA (*ssr*) and the RNA component of RNase P (*rnpB*) were designed by identifying reverse complement primers 20-30 bases long (optimum 25) with a T_m_ of 60-72°C (optimum 65°C) using *primer3*. Promiscuity to the remainder of the genome was checked by local *blastn*. To form the hairpin that guides Taq endonuclease cleavage (Figure S1C), oligo sequences were BINDINGREGION-<u>CATGCG</u>CTTG<u>CGCATG-</u>N, the underlined components indicate the predicted hairpin, N was the final base of the binding region ensuring a single base overhang^85^.

### Multiplex Taq-depleted Bacterial (MTB) RNA-Seq libraries

In 96 well plates, de-crosslinked RNA was back diluted in T.1E (10 mM Tris 8, 0.1 mM EDTA) to add 6 or 15 ng (isolate libraries) or 10, 50, or 100 ng (validation libraries) across 5 uL. RNA was mixed with premixed 2 uL 5x Maxima H-buffer and 0.25 uL 100 uM inline barcoded reverse transcription primer and incubated for 5 minutes at 95°C to fragment RNA and anneal primer. Then, a master mix of 0.5 uL 10 mM dNTPs, 1.75 uL water, 0.25 uL RNase inhibitor, and 0.25 uL Maxima H-reverse transcriptase was added for a final volume of 10 uL per reaction. On a thermocycler, reactions were incubated for 10 minutes at 25°C, 30 minutes at 50°C, and 5 minutes at 85°C. Half of each reaction was mixed (other half saved at −20°C) in groups of up to 48 samples with unique inline barcodes. To degrade free primers, for every 20 uL of mixed sample, 1 uL of exonuclease I was added, and reactions were incubated for 20 minutes at 37°C and 20 minutes at 80°C.

To clean up, reactions were brought to a minimum volume of 100 uL with water and mixed 1:1 with cleanup beads (11g PEG 8000, 10 mL 5 M NaCl, 100 uL 0.5 M EDTA, 50 uL 10% IGEPAL, 250 uL sodium azide, 1 mL Sera-Mag Speed Beads). Cleanups were incubated 5 minutes at room temperature followed by 5 minutes on a magnetic rack to clarify. Supernatant was discarded and beads were washed twice with a minimum of 400 uL (or the cleanup reaction volume if higher) of 80% ethanol. After the last wash, tubes were spun for a moment to collect ethanol at the bottom of the tube and excess was removed on the magnetic rack. Beads were allowed to dry for 1 minute. Beads were resuspended in 10 uL of T.1E and incubated for 5 minutes off the rack before returning to the rack and saving the 10 uL supernatant.

To degrade excess RNA, 1.5 uL of RNase H buffer, 3 uL of water, and 0.5 uL of RNase H was added to each reaction and reactions were incubated for 30 minutes at 37°C; reactions can be frozen or placed on ice when done. While RNase H reaction was proceeding, the adapter and splint oligos were pre-annealed by mixing adapter at 10 uM and splint at 20 uM final concentrations in T.1E and incubating at 95°C for 30 seconds and gradually lowering temperature to room temperature on a thermocycler; pre-annealed adapter can be saved at −20°C. To the RNase H reaction, we added 5 uL 10x T4 ligase buffer, 20 uL 50% PEG 8000, 1 uL pre-annealed adapter/splint, 8 uL water, and 1 uL T4 ligase (high concentration). Ligations were incubated for 1 hour at 25°C. Reactions were cleaned up by adding 50 uL water and 80 uL cleanup beads and following above cleanup protocol but resuspending in 20 uL of T.1E.

rRNA depletions were conducted by mixing 10 uL cDNA library from the prior step, 3 uL 10x Taq polymerase buffer, 2 uL Taq polymerase, water, and depletion primers. Depletion primers and water quantity was calculated to achieve a 0.1 uM final concentration of an individual primer. Reactions were incubated at 95°C for 30 seconds, 65°C for 2 hours, and held at 10°C. To clean up, reactions were brought to a 100 uL volume and cleaned up as above, resuspending in 20 uL of T.1E.

For each pool, a trial qPCR reaction was prepared to determine the correct number of final amplification cycles: 2 uL of cDNA library was mixed with 5 uL 2x iTaq master mix, 0.2 uL of each 100 uM Illumina adapter primer, and 2.6 uL of water. The qPCR machine was set to 5 minutes at 95°C, followed by 25 cycles of 30 seconds at 95°C and 45 seconds at 60°C. For each pool, the number of cycles where amplification was one quarter of the way through the sigmoidal amplification curve was recorded as the cycle number for the final PCR (16-19 cycles for our libraries).

Final PCRs were prepared as 50 uL reactions with 25 uL 2x KAPA HiFi master mix, 1 uL of each 100 uM Illumina adapter primer, 15 uL cDNA library and 8 uL of water; i7 indexes were varied to enable pooling of all sets of inline barcodes after PCR. The final PCR was run on a thermocycler set to 45 seconds at 98°C; followed by X cycles of 15 seconds at 98°C, 30 seconds at 60°C, 30 seconds at 72°C; followed by a final incubation of 1 minute at 72°C. “X” denotes the final cycle number determined by the trial qPCR.

A two-sided magnetic bead size selection/cleanup was conducted using a 20% PEG-NaCl solution (20% w/v PEG 8000, 2.5 M NaCl). PCRs were brought to a volume of 200 uL with water. For the first (upper) size selection, 4 uL of Sera-Mag Speed Beads were resuspended in 100 uL 20% PEG-NaCl and mixed with reaction and incubated 5 minutes at room temperature. Bead-reaction mixture was placed on a magnetic rack and allowed to clarify; the supernatant was transferred to a fresh tube and beads were discarded. For the second (lower) size selection, 4 uL Sera-Mag Speed Beads were resuspended in 40 uL 20% PEG-NaCl and mixed with the supernatant of the upper size selection and incubated 5 minutes at room temperature. The mixture was placed on a magnetic rack and allowed to clarify before removing and discarding supernatant. The beads were washed twice with 400 uL 80% ethanol while on the magnetic rack. After the last wash, tubes were spun for a moment to collect ethanol at the bottom of the tube and excess was removed on the magnetic rack. Beads were allowed to dry for 1 minute. Beads were resuspended in 10 uL of T.1E and incubated for 5 minutes off the rack before returning to the rack and saving the 10 uL library containing supernatant.

Library concentrations were quantified by qubit for pooling. Method validation sequencing was conducted on a MiSeq instrument. Quality control sequencing for isolate pools was conducted on a MiSeq instrument and library ratios were determined by sequencing to re-pool reverse transcription reactions at an ideal ratio. Final clinical isolate sequence pools were submitted for sequencing on a NovaSeq instrument to the Harvard Bauer Core Facility. Inline demultiplexing, mapping to the H37Rv (NC_000962.3) genome, and quality control was conducted with a pipeline built on *cutadapt* and *bowtie2*. Fragments were deduplicated with *picard*. Reads were filtered to remove ambiguously mapping reads to only include those with a *mapq* value greater than 10. Library depth normalization was conducted by calculating size factors relative to a pseudo-reference sample as in *deseq2*^86^. For validation libraries (Figures 1D-E), the top 50% of genes by mean expression across samples were considered well-expressed. For isolate libraries (Figures 2-6), a log_2_ normalized mean expression threshold of 3 was considered well-expressed to minimize standard deviation between replicates while keeping >50% of coding regions (Figure S2C).

### Analyses of strain gene expression diversity

To identify high expression diversity genes in Figure 2A and Table S2, we (1) identified genes in the top 5% of biologic (standard deviation of strain means) to technical variation (mean standard deviation of strain replicates) within well-expressed genes and (2) identified genes in the top 50% of absolute variation (standard deviation of strain means) within well-expressed genes. We conducted this analysis for all strains and for each of lineages 1, 2, and 4 separately. A gene was considered high expression diversity if it met both thresholds in across all strains or in at least 1 lineage. The results of the “all strains” analysis is shown in Figure 2A. For Figure 2B, the log_2_ normalized gene expression level relative to the mean for that gene is shown in phylogenetic order on the x-axis and by Ward clustering of high variation gene expression data (dendrogram not shown) on the y-axis. To generate Figure 2D, Ward clustering of high variation gene expression data was conducted separately on all strains and on each of lineages 1, 2, and 4. All pairwise distances were calculated from the Ward dendrogram (phenotypic distance) and the phylogenetic tree (phylogenetic distance) and normalized to the maximum distance. The Mantel test and R^2^ was used to describe the relationship between the two distance matrices.

Gene category enrichments in Figure 3A were calculated on the set of well-expressed genes searching for enrichment of expression diverse genes. Two-sided enrichment was calculated with the Fisher exact test against each *mycobrowser* category, genes previously identified as *in vitro* essential^34^, genes previously identified as *in vivo* essential across 30 collaborative cross mouse strains^35^, genes identified as containing an epitope across at least 2 references on the Immune Epitope Database^36^, and genes used as vaccine antigens identified by a literature search^37–43^.

### GWAS of gene expression and coding variants

GWAS in Mtb is complicated by clonal inheritance and a lack of horizontal gene transfer. This complicates GWAS by (1) introducing strong population biases due to complete linkage and (2) biasing acquisition of beneficial alleles towards *de novo* mutation. Particularly for selection towards loss of function at a locus, the requirement for *de novo* variation can mean many variants across a population can achieve the same phenotype but be present at a relatively low population frequency. To control for this low population frequency, we used burden testing to consider all predicted functional variants in each coding region as one variant class. Here, we define predicted functional variants as those predicted by *sift4g* to be deleterious (sift score < 0.05) or annotated as causing a frameshift, creating an early stop codon, or disrupting the start codon. To focus on modern selection and avoid associations driven by lineage-defining variants, we considered only variants that occur in less than 25% of strains and required loci to have at least 5 independent mutational events. We further removed potential low quality variant calls by removing variants with missing data in >5% of strains or had reversions (which should be very rare). Only variants called as Alt relative to H37Rv were considered. GWAS was conducted with the *pyseer* package using a linear mixed model and providing a strain similarity matrix built from the phylogenetic tree to control for population effects. The association was run iteratively against each of the 161 high expression diversity genes as a phenotype to generate 49,427 gene expression to variant loci associations. The reported FDR was calculated by Benjamini-Hochberg multiple hypothesis correction across all associations (Figure 3B, Table S3).

### SlfR protein purification and IDAP-Seq

SlfR protein purification and IDAP-Seq was adjusted from previously described^44^. *slfR* with an N-terminal 6xHIS tag was amplified from *slfR* from H37Rv genomic DNA inserted into pET-28 using the NcoI and HindIII sites and Gibson assembly. The resulting plasmid was isolated from NEB 5-alpha Competent E. coli and transformed into BL21-CodonPlus (DE3)-RP Competent Cells. The resulting strain was cultured overnight in LB with 50 µg/mL kanamycin with shaking at 37°C. The overnight culture was diluted in two 50 mL cultures of LB with kanamycin, grown up to OD600 = 0.5, and protein production was induced with 500 µM IPTG for 4 hours shaking at 37°C. Cells were pelleted at 5000 g for 15 minutes and frozen at −80°C. Pellets were resuspended in 10 mL binding buffer (500 mM Tris-HCl pH 8.0, 100 mM NaCl, 50 mM KCl, 3% glycerol) with protease inhibitor and lysozyme and lysed via sonication 4 x 30 seconds 30% amplitude, alternating with cooling on ice. Lysate was spun at 20,000 g at 4°C for 15 minutes and clarified protein lysate was removed and combined with 1 mL Ni-NTA Agarose and 10 mM imidazole and incubated in the column at 4°C for 1 hour, gently shaking. Supernatant was drained from the gravity flow column and agarose was washed twice with binding buffer with 20 mM imidazole. Protein was eluted with binding buffer with 100 mM imidazole (elution 1) followed by binding buffer with 200 mM imidazole (elution 2). Protein from elution 2 was dialyzed using a Slide-A-Lyzer G3 Dialysis Cassette (10K MWCO, 3 mL) in 200 mL dialysis buffer (50 mM Tris-HCl pH 8.0, 50 mM NaCl, 0.25 mg/mL BSA, 5% glycerol) for 1 hour at room temperature and then in a fresh 200 mL dialysis buffer overnight at 4°C. Resulting protein was measured by Qubit Protein Assay, aliquoted, flash frozen in liquid nitrogen, and stored at −80°C.

We prepared two replicates of each of 0 nM or 300 nM SlfR in dialysis buffer with 1 μg of sheared Mtb genomic DNA (mean 300 bp size) in a 125 uL reaction volume. Binding reactions were incubated 30 minutes at room temperature. For each reaction, 100 uL Talon resin was pre-equilibrated in dialysis buffer by centrifuging for 2 minutes at 700 g to remove supernatant and mixing with 500 uL dialysis buffer. Equilibration with dialysis buffer was repeated and supernatant was discarded. Next, reactions were mixed with a pre-equilibrated Talon resin and incubated 30 minutes at room temperature with rotation. Bound DnaA-resin mixtures were then loaded onto Poly-Prep chromatography columns and washed 3 times with 1 mL of dialysis buffer. Bound genomic DNA was eluted by capping the columns and adding 500 μL of pre-warmed (65°C) elution buffer (50 mM Tris pH 8, 10 mM EDTA, 1% SDS) and incubating columns on a heat block at 65°C for 15 minutes with the tops coated with parafilm to prevent evaporation. After incubation, elution buffer flow through was collected with two additional 200 uL washes of elution buffer. Eluted gDNA was brought to a 1 mL volume with additional elution buffer. Reactions were mixed with 10 uL of Sera-Mag Speed Beads and 1 mL of 20% PEG-NaCl (20% w/v PEG 8000, 2.5 M NaCl) and incubated for 5 minutes at room temperature. Beads were washed twice with 2 mL of 80% ethanol. After the last wash, tubes were spun for a moment to collect ethanol at the bottom of the tube and excess was removed on the magnetic rack. Beads were allowed to dry for 1 minute. Beads were resuspended in 20 uL of T.1E and incubated for 5 minutes off the rack before returning to the rack and saving the 20 uL supernatant. Libraries were prepared using the NEBNext Ultra II DNA Library Prep Kit for Illumina following manufacturer guidelines for <5 ng of input DNA. Libraries were sequenced with paired ends on a MiSeq instrument.

To map the binding of SlfR to the Mtb genome, the fragment density was calculated at every position by mapping paired end reads with *bowtie2* and depth normalized by dividing densities by a size factor (sample counts / geometric mean of all sample counts). Chromosomal regions bound by SlfR were identified by dividing the mean fragment density of SlfR + samples by the mean of SlfR – samples. Peak positions were called using *find_peaks* in *scipy* with a minimum distance of 500 bases between peaks and the top 50 peaks and nearby genes are shown in Table S4. Motif calling was conducted using *meme* on the 200 bases surrounding each peak position with a signal ratio greater than 5 allowing zero or one occurrence per sequence and enforcing palindromes.

### Proteomics of *slfR* variant strains

The following strain pairs (differing by independent *slfR* variants in lineage 2; variant strains marked with *) were grown: g2g_030* and vci_269, vci_246* and vci_089, vci_569* and vci_035. Strains were grown in a 10 mL volume in 30 mL inkwells at 37°C x 100 RPM, in 7H9 media supplemented as above. Cultures were grown for 3 days to mid-log (0.35-0.67) then 1-2 mL were back diluted into triplicate 50 mL cultures and grown for 4 additional days to mid-log (0.32-0.72). To remove detergents that can affect mass spectrometry results, cultures were centrifuged at 4,000 RPM for 7 minutes in a swinging bucket centrifuge to pellet cells and washed twice with 20 mL of 1x PBS before resuspending in 1 mL PBS. Cells were pelleted a final time on a benchtop centrifuge (10,000 RPM for 5 minutes) before freezing pellets at −80°C. Pellets were resuspended in 1 mL lysis buffer provided by the Whitehead Institute Proteomics facility (5% SDS, 100 mM TEAB, 40 mM CAA, 10 mM TCEP) and transferred to a bead beating tube. Cells were bead beaten 3 times for 30 seconds at maximum speed, resting on ice between steps. Bead beating tubes were spun 10 minutes at max speed on a benchtop centrifuge to remove debris and supernatant was placed in a fresh tube. Supernatant was filter sterilized twice through 0.22 μm syringe filters and submitted to Whitehead Institute Proteomics facility for DIA label-free full proteome analysis normalized across samples^87^. We took the mean of replicate sample protein abundances, masking any proteins not found in all samples, and then used a z-test without assuming equal variance to compare the difference between *slfR* variant and ancestral isolate abundances. The reported FDR was calculated by Benjamini-Hochberg multiple hypothesis correction across z-tests of individual protein abundance changes.

### Western blotting of *whiB6* variant strains

The following strain pairs (differing by independent *whiB6* variants in lineage 2, variant strains marked with *) were grown: vci_149* and vci_177, vci_176* and vci_403, vci_439* and vci_181. Strains were grown in a 10 mL volume in 30 mL inkwells at 37°C x 100 RPM, in 7H9 media supplemented as above. Strains were grown for 3 days to mid log (OD 0.49-0.61), then 1-2 mL were back diluted into duplicate 50 mL cultures in the same media and grown for 4 additional days to allow all cultures to get above OD 0.5. To prevent detergent effects on secretion, samples were then transferred to detergent-free Sauton’s media (for 1 L: 0.5 g KH_2_PO_4_, 0.5 g MgSO_4_·7H_2_O, 2 g citric acid, 0.05 g ferric ammonium citrate, 60 mL glycerol, 4 g asparagine; pH adjusted to 7.2 with NaOH) by pelleting cultures at 4,000 RPM for 7 minutes and washing twice with 20 mL Sauton’s media and normalizing all cultures to 0.5 OD in 50 mL of Sauton’s based on pre-spin OD. Cultures were grown for 24 hours then by pelleting at 4,000 RPM for 15 minutes. 45 mL of supernatant was filter sterilized twice through a 0.22 μm syringe filter. Pellets were resuspended in 1 mL protein extraction buffer (50 mM Tris HCl pH 7.5, 5 mM EDTA, 1x Roche cOmplete protease inhibitor) and bead beaten and sterilized as above pellets for proteomics.

The concentration of protein extracted from pellet was measured using Qubit Protein Assay. Higher protein concentration samples were diluted in water down to the lowest protein concentration then all samples were diluted 1:10 in water. Equal volumes of supernatant samples were taken. Samples were then incubated for 10 minutes at 95°C in 6X Laemmli SDS-Sample Buffer. SeeBlue™ Plus2 Pre-stained Protein Standard and 18µL of treated samples were then loaded onto NuPAGE™ 4 to 12%, Bis-Tris gels and run in NuPAGE MES SDS Running Buffer for 30-45 minutes at 170 V. Protein was transferred onto nitrocellulose membranes using 10X Transfer Buffer diluted in methanol and water (1:1:8), on a Trans-Blot Turbo Transfer System (BioRad) for 30 minutes at 25 V and 1 A. Membranes were rinsed with 1x PBS and blocked for two hours at room temperature using Blocking Buffer for Fluorescent Western Blotting. Primary antibody (antibodies to CFP-10, GroEL2, and Mpt64) were diluted in 1:1000 (CFP-10, GroEL2) or 1:5000 (Mpt64) in blocking buffer supplemented with 0.1% Tween-20. Membranes were incubated in primary antibody overnight at 4°C then washed with PBS with 0.05% Tween-20. LI-COR Secondary antibodies were diluted 1:20000 in blocking buffer with 0.1% Tween-20 and 0.0025% SDS. Membranes were incubated in relevant secondary antibody for 1 hour at room temperature, washed with PBS with 0.05% Tween-20, and left in PBS before imaging. Images were acquired with LI-COR Odyssey CLx Imager. Blot quantifications were done in EmpiriaStudio using automatic lane finder and background adjustments and manually adjusted band finding. Display adjustments for figure visualizations were also done in EmpiriaStudio.

### Bioinformatic analysis details for *whiB6* and *espACD* upstream variants

As with above GWAS predicted functional variants, *whiB6* predicted functional variants were defined as those predicted by *sift4g* to be deleterious (sift score < 0.05) or annotated as causing a frameshift, creating an early stop codon, or disrupting the start codon. To generate cumulative variant strain plots, we used our parsimony ancestral reconstructions to identify ancestral nodes where *whiB6* variants occurred, ordered them by the number of strains inheriting the variant and calculated the cumulative number of *whiB6* variant strains across each individual lineage.

For cumulative distribution of *espA* upstream variants, only deletions exactly matching del A and B are shown. Upstream variant sites were defined as the positions of sites A and B (4057190 and 4057367) ±5 bases. Cumulative distributions were plotted as described above.

### Bioinformatic analysis details for antibiotic resistance associations, transmission, and timing

To identify strains with high-level drug resistance (DR), we used the *tbprofiler* database of genotypic DR. We classified strains with one or more DR variants “Associated with Resistance” or “Associated with Resistance – Interim” in the WHO catalogue v2 as DR to the corresponding drug. To identify associations between predicted functional coding variants in *whiB6* or *slfR* (defined as above) or variants upstream of *espACD* (both deletions and site variants as defined above), we applied a recoded version of the *phyoverlap* algorithm designed to run on larger trees^68^. Briefly, for each independent mutational event of a target mutation (e.g. predicted functional *whiB6* variants) identified by parsimony, we calculated the fraction of strains inheriting that variant that had genotypic DR as defined above, calculating an overlap fraction. Next, we took the mean of overlap fractions for the target variants to calculate an observed overlap statistic. We compared the overlap statistic to control values by randomly redistributing the observed number of target mutational events across the tree (where the probability of selecting a branch was proportional to its length on the phylogenetic tree) and recalculating the overlap statistic. To generate a null distribution, we repeated this up to 10,000 times; the estimated p-value was the number of permutations greater than the observed overlap statistic divided by the number of permutations run. This analysis was performed separately on each lineage’s subtree.

We used a similar algorithm to identify associations between variants and transmission or modernity. In this case, the mean terminal branch length (transmission) or branch midpoint’s distance from root (modernity) was pre-calculated for every node on the phylogenetic tree. Then, for each independent mutational event of the target mutation identified by parsimony, the node’s value was pulled from precalculated values; the mean of all values was taken to calculate an observed statistic. Null distributions were calculated by permutations as above. As transmission and modernity are two-sided—i.e. we were interested in variants associated with terminal branch lengths shorter or longer than the null distribution or variants more ancient or modern than the null distribution—we reported estimated p-values based on the smaller of permutations greater than or less than the observed statistic.

Associations for DS or DR strains were calculated using the same algorithm, but on trees with or without any resistance mutations as defined above. To generate these trees, we started with the full tree and pruned all strains with or without resistance using *dendropy*. Ancestral reconstruction was then re-run on the DS and DR trees.

## References

1. Boritsch, E.C., Khanna, V., Pawlik, A., Honoré, N., Navas, V.H., Ma, L., Bouchier, C., Seemann, T., Supply, P., Stinear, T.P., et al. (2016). Key experimental evidence of chromosomal DNA transfer among selected tuberculosis-causing mycobacteria. Proc. Natl. Acad. Sci. 113, 9876–9881. 10.1073/pnas.1604921113.

2. Didelot, X., Bowden, R., Wilson, D.J., Peto, T.E.A., and Crook, D.W. (2012). Transforming clinical microbiology with bacterial genome sequencing. Nat. Rev. Genet. 13, 601–612. 10.1038/nrg3226.

3. Bastos, H.N., Osório, N.S., Gagneux, S., Comas, I., and Saraiva, M. (2018). The Troika Host– Pathogen–Extrinsic Factors in Tuberculosis: Modulating Inflammation and Clinical Outcomes. Front Immunol 8, 1948. 10.3389/fimmu.2017.01948.

4. Goig, G.A., Windels, E.M., Loiseau, C., Stritt, C., Biru, L., Borrell, S., Brites, D., and Gagneux, S. (2025). Ecology, global diversity and evolutionary mechanisms in the Mycobacterium tuberculosis complex. Nat. Rev. Microbiol., 1–13. 10.1038/s41579-025-01159-w.

5. Liu, Q., Zhu, J., Dulberger, C.L., Stanley, S., Wilson, S., Chung, E.S., Wang, X., Culviner, P., Liu, Y.J., Hicks, N.D., et al. (2022). Tuberculosis treatment failure associated with evolution of antibiotic resilience. Science 378, 1111–1118. 10.1126/science.abq2787.

6. Chiner-Oms, Á., López, M.G., Moreno-Molina, M., Furió, V., and Comas, I. (2022). Gene evolutionary trajectories in Mycobacterium tuberculosis reveal temporal signs of selection. Proc. Natl. Acad. Sci. 119, e2113600119. 10.1073/pnas.2113600119.

7. Wray, G.A., Hahn, M.W., Abouheif, E., Balhoff, J.P., Pizer, M., Rockman, M.V., and Romano, L.A. (2003). The Evolution of Transcriptional Regulation in Eukaryotes. Mol. Biol. Evol. 20, 1377–1419. 10.1093/molbev/msg140.

8. Babu, M.M., Teichmann, S.A., and Aravind, L. (2006). Evolutionary Dynamics of Prokaryotic Transcriptional Regulatory Networks. J. Mol. Biol. 358, 614–633. 10.1016/j.jmb.2006.02.019.

9. Gomez-Gonzalez, P.J., Andreu, N., Phelan, J.E., Sessions, P.F. de, Glynn, J.R., Crampin, A.C., Campino, S., Butcher, P.D., Hibberd, M.L., and Clark, T.G. (2019). An integrated whole genome analysis of Mycobacterium tuberculosis reveals insights into relationship between its genome, transcriptome and methylome. Sci Rep-uk 9, 5204. 10.1038/s41598-019-41692-2.

10. Chiner-Oms, Á., Berney, M., Boinett, C., González-Candelas, F., Young, D.B., Gagneux, S., Jacobs, W.R., Parkhill, J., Cortes, T., and Comas, I. (2019). Genome-wide mutational biases fuel transcriptional diversity in the Mycobacterium tuberculosis complex. Nat Commun 10, 3994. 10.1038/s41467-019-11948-6.

11. Holt, K.E., McAdam, P., Thai, P.V.K., Thuong, N.T.T., Ha, D.T.M., Lan, N.N., Lan, N.H., Nhu, N.T.Q., Hai, H.T., Ha, V.T.N., et al. (2018). Frequent transmission of the Mycobacterium tuberculosis Beijing lineage and positive selection for the EsxW Beijing variant in Vietnam. Nat Genet 50, 849–856. 10.1038/s41588-018-0117-9.

12. Luo, Y., Huang, C.-C., Howard, N.C., Wang, X., Liu, Q., Li, X., Zhu, J., Amariuta, T., Asgari, S., Ishigaki, K., et al. (2024). Paired analysis of host and pathogen genomes identifies determinants of human tuberculosis. Nat. Commun. 15, 10393. 10.1038/s41467-024-54741-w.

13. Coscolla, M., Copin, R., Sutherland, J., Gehre, F., de Jong, B., Owolabi, O., Mbayo, G., Giardina, F., Ernst, J.D., and Gagneux, S. (2015). M. tuberculosis T Cell Epitope Analysis Reveals Paucity of Antigenic Variation and Identifies Rare Variable TB Antigens. Cell Host Microbe 18, 538–548. 10.1016/j.chom.2015.10.008.

14. Pym, A.S., Brodin, P., Brosch, R., Huerre, M., and Cole, S.T. (2002). Loss of RD1 contributed to the attenuation of the live tuberculosis vaccines Mycobacterium bovis BCG and Mycobacterium microti. Mol Microbiol 46, 709–717. 10.1046/j.1365-2958.2002.03237.x.

15. Anderson, R.M., and May, R.M. (1982). Coevolution of hosts and parasites. Parasitology 85, 411–426. 10.1017/s0031182000055360.

16. Barrick, J.E., Colburn, G., Deatherage, D.E., Traverse, C.C., Strand, M.D., Borges, J.J., Knoester, D.B., Reba, A., and Meyer, A.G. (2014). Identifying structural variation in haploid microbial genomes from short-read resequencing data using breseq. Bmc Genomics 15, 1039. 10.1186/1471-2164-15-1039.

17. Phelan, J.E., O’Sullivan, D.M., Machado, D., Ramos, J., Oppong, Y.E.A., Campino, S., O’Grady, J., McNerney, R., Hibberd, M.L., Viveiros, M., et al. (2019). Integrating informatics tools and portable sequencing technology for rapid detection of resistance to anti-tuberculous drugs. Genome Med. 11, 41. 10.1186/s13073-019-0650-x.

18. Danecek, P., Bonfield, J.K., Liddle, J., Marshall, J., Ohan, V., Pollard, M.O., Whitwham, A., Keane, T., McCarthy, S.A., Davies, R.M., et al. (2021). Twelve years of SAMtools and BCFtools. GigaScience 10, giab008. 10.1093/gigascience/giab008.

19. Li, H., and Durbin, R. (2009). Fast and accurate short read alignment with Burrows–Wheeler transform. Bioinformatics 25, 1754–1760. 10.1093/bioinformatics/btp324.

20. McKenna, A., Hanna, M., Banks, E., Sivachenko, A., Cibulskis, K., Kernytsky, A., Garimella, K., Altshuler, D., Gabriel, S., Daly, M., et al. (2010). The Genome Analysis Toolkit: A MapReduce framework for analyzing next-generation DNA sequencing data. Genome Res. 20, 1297–1303. 10.1101/gr.107524.110.

21. Ishikawa, S.A., Zhukova, A., Iwasaki, W., and Gascuel, O. (2019). A Fast Likelihood Method to Reconstruct and Visualize Ancestral Scenarios. Mol. Biol. Evol. 36, 2069–2085. 10.1093/molbev/msz131.

22. Price, M.N., Dehal, P.S., and Arkin, A.P. (2010). FastTree 2 – Approximately Maximum-Likelihood Trees for Large Alignments. Plos One 5, e9490. 10.1371/journal.pone.0009490.

23. Vargas, R., Luna, M.J., Freschi, L., Marin, M., Froom, R., Murphy, K.C., Campbell, E.A., Ioerger, T.R., Sassetti, C.M., and Farhat, M.R. (2023). Phase variation as a major mechanism of adaptation in Mycobacterium tuberculosis complex. Proc. Natl. Acad. Sci. 120, e2301394120. 10.1073/pnas.2301394120.

24. Schloissnig, S., Arumugam, M., Sunagawa, S., Mitreva, M., Tap, J., Zhu, A., Waller, A., Mende, D.R., Kultima, J.R., Martin, J., et al. (2013). Genomic variation landscape of the human gut microbiome. Nature 493, 45–50. 10.1038/nature11711.

25. Turkarslan, S., Peterson, E.J.R., Rustad, T.R., Minch, K.J., Reiss, D.J., Morrison, R., Ma, S., Price, N.D., Sherman, D.R., and Baliga, N.S. (2015). A comprehensive map of genome-wide gene regulation in Mycobacterium tuberculosis. Sci. Data 2, 150010. 10.1038/sdata.2015.10.

26. Kapopoulou, A., Lew, J.M., and Cole, S.T. (2011). The MycoBrowser portal: A comprehensive and manually annotated resource for mycobacterial genomes. Tuberculosis 91, 8–13. 10.1016/j.tube.2010.09.006.

27. Kachroo, P., Eraso, J.M., Beres, S.B., Olsen, R.J., Zhu, L., Nasser, W., Bernard, P.E., Cantu, C.C., Saavedra, M.O., Arredondo, M.J., et al. (2019). Integrated analysis of population genomics, transcriptomics and virulence provides novel insights into Streptococcus pyogenes pathogenesis. Nat. Genet. 51, 548–559. 10.1038/s41588-018-0343-1.

28. Lyamichev, V., Brow, M., and Dahlberg, J. (1993). Structure-specific endonucleolytic cleavage of nucleic acids by eubacterial DNA polymerases. Science 260, 778–783. 10.1126/science.7683443.

29. Gu, W., Crawford, E.D., O’Donovan, B.D., Wilson, M.R., Chow, E.D., Retallack, H., and DeRisi, J.L. (2016). Depletion of Abundant Sequences by Hybridization (DASH): using Cas9 to remove unwanted high-abundance species in sequencing libraries and molecular counting applications. Genome Biol 17, 41. 10.1186/s13059-016-0904-5.

30. Mehra, S., Foreman, T.W., Didier, P.J., Ahsan, M.H., Hudock, T.A., Kissee, R., Golden, N.A., Gautam, U.S., Johnson, A.-M., Alvarez, X., et al. (2015). The DosR Regulon Modulates Adaptive Immunity and Is Essential for Mycobacterium tuberculosis Persistence. Am J Resp Crit Care 191, 1185–1196. 10.1164/rccm.201408-1502oc.

31. McKinney, J.D., Bentrup, K.H. zu, Muñoz-Elías, E.J., Miczak, A., Chen, B., Chan, W.-T., Swenson, D., Sacchettini, J.C., Jacobs, W.R., and Russell, D.G. (2000). Persistence of Mycobacterium tuberculosis in macrophages and mice requires the glyoxylate shunt enzyme isocitrate lyase. Nature 406, 735–738. 10.1038/35021074.

32. Belardinelli, J.M., Larrouy-Maumus, G., Jones, V., Carvalho, L.P.S. de, McNeil, M.R., and Jackson, M. (2014). Biosynthesis and Translocation of Unsulfated Acyltrehaloses in Mycobacterium tuberculosis *. J. Biol. Chem. 289, 27952–27965. 10.1074/jbc.m114.581199.

33. Resstel, C., Madduri, B.T.S.A., and Bell, S.L. (2024). Mycobacterial PE/PPE proteins function as “personal protective equipment” against host defenses. Front. Tuberc. 2, 1458105. 10.3389/ftubr.2024.1458105.

34. DeJesus, M.A., Gerrick, E.R., Xu, W., Park, S.W., Long, J.E., Boutte, C.C., Rubin, E.J., Schnappinger, D., Ehrt, S., Fortune, S.M., et al. (2017). Comprehensive Essentiality Analysis of the Mycobacterium tuberculosis Genome via Saturating Transposon Mutagenesis. Mbio 8, e02133–16. 10.1128/mbio.02133-16.

35. Smith, C.M., Baker, R.E., Proulx, M.K., Mishra, B.B., Long, J.E., Park, S.W., Lee, H.-N., Kiritsy, M.C., Bellerose, M.M., Olive, A.J., et al. (2022). Host-pathogen genetic interactions underlie tuberculosis susceptibility in genetically diverse mice. Elife 11, e74419. 10.7554/elife.74419.

36. Vita, R., Mahajan, S., Overton, J.A., Dhanda, S.K., Martini, S., Cantrell, J.R., Wheeler, D.K., Sette, A., and Peters, B. (2019). The Immune Epitope Database (IEDB): 2018 update. Nucleic Acids Res. 47, D339–D343. 10.1093/nar/gky1006.

37. Kaufmann, S.H.E. (2023). Vaccine development against tuberculosis before and after Covid-19. Front. Immunol. 14, 1273938. 10.3389/fimmu.2023.1273938.

38. Woodworth, J.S., Clemmensen, H.S., Battey, H., Dijkman, K., Lindenstrøm, T., Laureano, R.S., Taplitz, R., Morgan, J., Aagaard, C., Rosenkrands, I., et al. (2021). A Mycobacterium tuberculosis-specific subunit vaccine that provides synergistic immunity upon co-administration with Bacillus Calmette-Guérin. Nat. Commun. 12, 6658. 10.1038/s41467-021-26934-0.

39. Sergeeva, M., Romanovskaya-Romanko, E., Zabolotnyh, N., Pulkina, A., Vasilyev, K., Shurigina, A.P., Buzitskaya, J., Zabrodskaya, Y., Fadeev, A., Vasin, A., et al. (2021). Mucosal Influenza Vector Vaccine Carrying TB10.4 and HspX Antigens Provides Protection against Mycobacterium tuberculosis in Mice and Guinea Pigs. Vaccines 9, 394. 10.3390/vaccines9040394.

40. Chen, L., Xu, M., Wang, Z., Chen, B., Du, W., Su, C., Shen, X., Zhao, A., Dong, N., Wang, Y., et al. (2010). The development and preliminary evaluation of a new Mycobacterium tuberculosis vaccine comprising Ag85b, HspX and CFP-10:ESAT-6 fusion protein with CpG DNA and aluminum hydroxide adjuvants. FEMS Immunol. Méd. Microbiol. 59, 42–52. 10.1111/j.1574-695x.2010.00660.x.

41. Wilkie, M., Satti, I., Minhinnick, A., Harris, S., Riste, M., Ramon, R.L., Sheehan, S., Thomas, Z.-R.M., Wright, D., Stockdale, L., et al. (2020). A phase I trial evaluating the safety and immunogenicity of a candidate tuberculosis vaccination regimen, ChAdOx1 85A prime – MVA85A boost in healthy UK adults. Vaccine 38, 779–789. 10.1016/j.vaccine.2019.10.102.

42. Tait, D.R., Hatherill, M., Meeren, O.V.D., Ginsberg, A.M., Brakel, E.V., Salaun, B., Scriba, T.J., Akite, E.J., Ayles, H.M., Bollaerts, A., et al. (2019). Final Analysis of a Trial of M72/AS01E Vaccine to Prevent Tuberculosis. N. Engl. J. Med. 381, 2429–2439. 10.1056/nejmoa1909953.

43. Rais, M., Abdelaal, H., Reese, V.A., Ferede, D., Larsen, S.E., Pecor, T., Erasmus, J.H., Archer, J., Khandhar, A.P., Cooper, S.K., et al. (2023). Immunogenicity and protection against Mycobacterium avium with a heterologous RNA prime and protein boost vaccine regimen. Tuberculosis 138, 102302. 10.1016/j.tube.2022.102302.

44. Hicks, N.D., Giffen, S.R., Culviner, P.H., Chao, M.C., Dulberger, C.L., Liu, Q., Stanley, S., Brown, J., Sixsmith, J., Wolf, I.D., et al. (2020). Mutations in dnaA and a cryptic interaction site increase drug resistance in Mycobacterium tuberculosis. Plos Pathog 16, e1009063. 10.1371/journal.ppat.1009063.

45. Lees, J.A., Galardini, M., Bentley, S.D., Weiser, J.N., and Corander, J. (2018). pyseer: a comprehensive tool for microbial pangenome-wide association studies. Bioinform Oxf Engl 34, 4310–4312. 10.1093/bioinformatics/bty539.

46. Martini, M.C., Hicks, N.D., Xiao, J., Alonso, M.N., Barbier, T., Sixsmith, J., Fortune, S.M., and Shell, S.S. (2022). Loss of RNase J leads to multi-drug tolerance and accumulation of highly structured mRNA fragments in Mycobacterium tuberculosis. PLoS Pathog. 18, e1010705. 10.1371/journal.ppat.1010705.

47. Converse, S.E., Mougous, J.D., Leavell, M.D., Leary, J.A., Bertozzi, C.R., and Cox, J.S. (2003). MmpL8 is required for sulfolipid-1 biosynthesis and Mycobacterium tuberculosis virulence. Proc. Natl. Acad. Sci. 100, 6121–6126. 10.1073/pnas.1030024100.

48. Domenech, P., Reed, M.B., Dowd, C.S., Manca, C., Kaplan, G., and Barry, C.E. (2004). The Role of MmpL8 in Sulfatide Biogenesis and Virulence of Mycobacterium tuberculosis *. J. Biol. Chem. 279, 21257–21265. 10.1074/jbc.m400324200.

49. Gangadharam, P.R.J., Cohn, M.L., and Middlebrook, G. (1963). Infectivity, pathogenicity and sulpholipid fraction of some Indian and British strains of tubercle bacilli. Tubercle 44, 452–455. 10.1016/s0041-3879(63)80087-2.

50. Ruhl, C.R., Pasko, B.L., Khan, H.S., Kindt, L.M., Stamm, C.E., Franco, L.H., Hsia, C.C., Zhou, M., Davis, C.R., Qin, T., et al. (2020). Mycobacterium tuberculosis Sulfolipid-1 Activates Nociceptive Neurons and Induces Cough. Cell 181, 293–305.e11. 10.1016/j.cell.2020.02.026.

51. Smith, J.L., and Grossman, A.D. (2015). In Vitro Whole Genome DNA Binding Analysis of the Bacterial Replication Initiator and Transcription Factor DnaA. Plos Genet 11, e1005258. 10.1371/journal.pgen.1005258.

52. Solans, L., Aguiló, N., Samper, S., Pawlik, A., Frigui, W., Martín, C., Brosch, R., and Gonzalo-Asensio, J. (2014). A Specific Polymorphism in Mycobacterium tuberculosis H37Rv Causes Differential ESAT-6 Expression and Identifies WhiB6 as a Novel ESX-1 Component. Infect Immun 82, 3446–3456. 10.1128/iai.01824-14.

53. Gröschel, M.I., Sayes, F., Simeone, R., Majlessi, L., and Brosch, R. (2016). ESX secretion systems: mycobacterial evolution to counter host immunity. Nat Rev Microbiol 14, 677–691. 10.1038/nrmicro.2016.131.

54. Stanley, S.A., Johndrow, J.E., Manzanillo, P., and Cox, J.S. (2007). The Type I IFN Response to Infection with Mycobacterium tuberculosis Requires ESX-1-Mediated Secretion and Contributes to Pathogenesis. J Immunol 178, 3143–3152. 10.4049/jimmunol.178.5.3143.

55. Simeone, R., Bobard, A., Lippmann, J., Bitter, W., Majlessi, L., Brosch, R., and Enninga, J. (2012). Phagosomal Rupture by Mycobacterium tuberculosis Results in Toxicity and Host Cell Death. Plos Pathog 8, e1002507. 10.1371/journal.ppat.1002507.

56. Koo, I.C., Wang, C., Raghavan, S., Morisaki, J.H., Cox, J.S., and Brown, E.J. (2008). ESX-1-dependent cytolysis in lysosome secretion and inflammasome activation during mycobacterial infection. Cell. Microbiol. 10, 1866–1878. 10.1111/j.1462-5822.2008.01177.x.

57. Damen, M.P.M., Meijers, A.S., Keizer, E.M., Piersma, S.R., Jiménez, C.R., Kuijl, C.P., Bitter, W., and Houben, E.N.G. (2022). The ESX-1 Substrate PPE68 Has a Key Function in ESX-1-Mediated Secretion in Mycobacterium marinum. mBio 13, e02819–22. 10.1128/mbio.02819-22.

58. Netikul, T., Palittapongarnpim, P., Thawornwattana, Y., and Plitphonganphim, S. (2021). Estimation of the global burden of Mycobacterium tuberculosis lineage 1. Infect., Genet. Evol. 91, 104802. 10.1016/j.meegid.2021.104802.

59. Fortune, S.M., Jaeger, A., Sarracino, D.A., Chase, M.R., Sassetti, C.M., Sherman, D.R., Bloom, B.R., and Rubin, E.J. (2005). Mutually dependent secretion of proteins required for mycobacterial virulence. Proc National Acad Sci 102, 10676–10681. 10.1073/pnas.0504922102.

60. Chen, J.M., Zhang, M., Rybniker, J., Basterra, L., Dhar, N., Tischler, A.D., Pojer, F., and Cole, S.T. (2013). Phenotypic Profiling of Mycobacterium tuberculosis EspA Point Mutants Reveals that Blockage of ESAT-6 and CFP-10 Secretion In Vitro Does Not Always Correlate with Attenuation of Virulence. J. Bacteriol. 195, 5421–5430. 10.1128/jb.00967-13.

61. Garces, A., Atmakuri, K., Chase, M.R., Woodworth, J.S., Krastins, B., Rothchild, A.C., Ramsdell, T.L., Lopez, M.F., Behar, S.M., Sarracino, D.A., et al. (2010). EspA Acts as a Critical Mediator of ESX1-Dependent Virulence in Mycobacterium tuberculosis by Affecting Bacterial Cell Wall Integrity. Plos Pathog 6, e1000957. 10.1371/journal.ppat.1000957.

62. Pang, X., Samten, B., Cao, G., Wang, X., Tvinnereim, A.R., Chen, X.-L., and Howard, S.T. (2013). MprAB Regulates the espA Operon in Mycobacterium tuberculosis and Modulates ESX-1 Function and Host Cytokine Response. J Bacteriol 195, 66–75. 10.1128/jb.01067-12.

63. Rosenberg, O.S., Dovey, C., Tempesta, M., Robbins, R.A., Finer-Moore, J.S., Stroud, R.M., and Cox, J.S. (2011). EspR, a key regulator of Mycobacterium tuberculosis virulence, adopts a unique dimeric structure among helix-turn-helix proteins. Proc. Natl. Acad. Sci. 108, 13450– 13455. 10.1073/pnas.1110242108.

64. Broset, E., Martín, C., and Gonzalo-Asensio, J. (2015). Evolutionary Landscape of the Mycobacterium tuberculosis Complex from the Viewpoint of PhoPR: Implications for Virulence Regulation and Application to Vaccine Development. mBio 6, 10.1128/mbio.01289-15. https://doi.org/10.1128/mbio.01289-15.

65. Luna, M.J., Oluoch, P.O., Miao, J., Culviner, P., Papavinasasundaram, K., Jaecklein, E., Shell, S.S., Ioerger, T.R., Fortune, S.M., Farhat, M.R., et al. (2025). Frequently arising ESX-1-associated phase variants influence Mycobacterium tuberculosis fitness in the presence of host and antibiotic pressures. mBio, e0376224. 10.1128/mbio.03762-24.

66. Global tuberculosis report (2024). (World Health Organization).

67. Levin-Reisman, I., Ronin, I., Gefen, O., Braniss, I., Shoresh, N., and Balaban, N.Q. (2017). Antibiotic tolerance facilitates the evolution of resistance. Science 355, 826–830. 10.1126/science.aaj2191.

68. Hicks, N.D., Yang, J., Zhang, X., Zhao, B., Grad, Y.H., Liu, L., Ou, X., Chang, Z., Xia, H., Zhou, Y., et al. (2018). Clinically prevalent mutations in Mycobacterium tuberculosis alter propionate metabolism and mediate multidrug tolerance. Nat Microbiol 3, 1032–1042. 10.1038/s41564-018-0218-3.

69. Menardo, F. (2022). Understanding drivers of phylogenetic clustering and terminal branch lengths distribution in epidemics of Mycobacterium tuberculosis. eLife 11, e76780. 10.7554/elife.76780.

70. Freschi, L., Vargas, R., Husain, A., Kamal, S.M.M., Skrahina, A., Tahseen, S., Ismail, N., Barbova, A., Niemann, S., Cirillo, D.M., et al. (2021). Population structure, biogeography and transmissibility of Mycobacterium tuberculosis. Nat. Commun. 12, 6099. 10.1038/s41467-021-26248-1.

71. Walter, K.S., Santos, P.C.P. dos, Gonçalves, T.O., Silva, B.O. da, Santos, A. da S., Leite, A. de C., Silva, A.M. da, Moreira, F.M.F., Oliveira, R.D. de, Lemos, E.F., et al. (2022). The role of prisons in disseminating tuberculosis in Brazil: A genomic epidemiology study. Lancet Reg. Heal. - Am. 9, 100186. 10.1016/j.lana.2022.100186.

72. Safi, H., Gopal, P., Lingaraju, S., Ma, S., Levine, C., Dartois, V., Yee, M., Li, L., Blanc, L., Liang, H.-P.H., et al. (2019). Phase variation in Mycobacterium tuberculosis glpK produces transiently heritable drug tolerance. Proc National Acad Sci 116, 19665–19674. 10.1073/pnas.1907631116.

73. Mayer-Barber, K.D., Andrade, B.B., Oland, S.D., Amaral, E.P., Barber, D.L., Gonzales, J., Derrick, S.C., Shi, R., Kumar, N.P., Wei, W., et al. (2014). Host-directed therapy of tuberculosis based on interleukin-1 and type I interferon crosstalk. Nature 511, 99–103. 10.1038/nature13489.

74. Manca, C., Tsenova, L., Bergtold, A., Freeman, S., Tovey, M., Musser, J.M., Barry, C.E., Freedman, V.H., and Kaplan, G. (2001). Virulence of a Mycobacterium tuberculosis clinical isolate in mice is determined by failure to induce Th1 type immunity and is associated with induction of IFN-α/β. Proc National Acad Sci 98, 5752–5757. 10.1073/pnas.091096998.

75. Chen, Z., Hu, Y., Cumming, B.M., Lu, P., Feng, L., Deng, J., Steyn, A.J.C., and Chen, S. (2016). Mycobacterial WhiB6 Differentially Regulates ESX-1 and the Dos Regulon to Modulate Granuloma Formation and Virulence in Zebrafish. Cell Reports 16, 2512–2524. 10.1016/j.celrep.2016.07.080.

76. Zwyer, M., Rutaihwa, L.K., Windels, E., Hella, J., Menardo, F., Sasamalo, M., Sommer, G., Schmülling, L., Borrell, S., Reinhard, M., et al. (2023). Back-to-Africa introductions of Mycobacterium tuberculosis as the main cause of tuberculosis in Dar es Salaam, Tanzania. PLOS Pathog. 19, e1010893. 10.1371/journal.ppat.1010893.

77. Jackson, M. (2014). The Mycobacterial Cell Envelope—Lipids. Cold Spring Harb. Perspect. Med. 4, a021105. 10.1101/cshperspect.a021105.

78. Arbues, A., Aguilo, J.I., Gonzalo-Asensio, J., Marinova, D., Uranga, S., Puentes, E., Fernandez, C., Parra, A., Cardona, P.J., Vilaplana, C., et al. (2013). Construction, characterization and preclinical evaluation of MTBVAC, the first live-attenuated M. tuberculosis-based vaccine to enter clinical trials. Vaccine 31, 4867–4873. 10.1016/j.vaccine.2013.07.051.

79. Jong, B.C. de, Hill, P.C., Brookes, R.H., Gagneux, S., Jeffries, D.J., Otu, J.K., Donkor, S.A., Fox, A., McAdam, K.P.W.J., Small, P.M., et al. (2006). Mycobacterium africanum Elicits an Attenuated T Cell Response to Early Secreted Antigenic Target, 6 kDa, in Patients with Tuberculosis and Their Household Contacts. J. Infect. Dis. 193, 1279–1286. 10.1086/502977.

80. Clemmensen, H.S., Knudsen, N.P.H., Rasmussen, E.M., Winkler, J., Rosenkrands, I., Ahmad, A., Lillebaek, T., Sherman, D.R., Andersen, P.L., and Aagaard, C. (2017). An attenuated Mycobacterium tuberculosis clinical strain with a defect in ESX-1 secretion induces minimal host immune responses and pathology. Sci. Rep. 7, 46666. 10.1038/srep46666.

81. Yoo, R., Rychel, K., Poudel, S., Al-bulushi, T., Yuan, Y., Chauhan, S., Lamoureux, C., Palsson, B.O., and Sastry, A. (2022). Machine Learning of All Mycobacterium tuberculosis H37Rv RNA-seq Data Reveals a Structured Interplay between Metabolism, Stress Response, and Infection. Msphere 7, e00033–22. 10.1128/msphere.00033-22.

82. Vaser, R., Adusumalli, S., Leng, S.N., Sikic, M., and Ng, P.C. (2016). SIFT missense predictions for genomes. Nat. Protoc. 11, 1–9. 10.1038/nprot.2015.123.

83. Cingolani, P., Platts, A., Wang, L.L., Coon, M., Nguyen, T., Wang, L., Land, S.J., Lu, X., and Ruden, D.M. (2012). A program for annotating and predicting the effects of single nucleotide polymorphisms, SnpEff. Fly 6, 80–92. 10.4161/fly.19695.

84. Nei, M., and Gojobori, T. (1986). Simple methods for estimating the numbers of synonymous and nonsynonymous nucleotide substitutions. Mol. Biol. Evol. 3, 418–426. 10.1093/oxfordjournals.molbev.a040410.

85. Nakano, M., Moody, E.M., Liang, J., and Bevilacqua, P.C. (2002). Selection for Thermodynamically Stable DNA Tetraloops Using Temperature Gradient Gel Electrophoresis Reveals Four Motifs: d(cGNNAg), d(cGNABg), d(cCNNGg), and d(gCNNGc) †. Biochemistry-us 41, 14281–14292. 10.1021/bi026479k.

86. Love, M.I., Huber, W., and Anders, S. (2014). Moderated estimation of fold change and dispersion for RNA-seq data with DESeq2. Genome Biol 15, 550. 10.1186/s13059-014-0550-8.

87. Demichev, V., Messner, C.B., Vernardis, S.I., Lilley, K.S., and Ralser, M. (2020). DIA-NN: Neural networks and interference correction enable deep proteome coverage in high throughput. Nat. methods 17, 41–44. 10.1038/s41592-019-0638-x.

